# A Site-Specific Organometallic Approach for Installing Tyrosine Phosphorylation Mimics to Decipher the Role of Phosphorylation in Alpha-Synuclein (aSyn) Aggregation and Seeding

**DOI:** 10.64898/2025.12.05.692498

**Authors:** Shaswati Mandal, Xiaoxi Lin, Anastasia V. Gemelli, Manjeet Singh, Anne-Laure M. Mellier, Yllza Jasiqi, Buchra Bouri, Muhammad Jbara, Hilal A. Lashuel

**Affiliations:** Laboratory of Molecular and Chemical Biology of Neurodegeneration, Institute of Bioengineering, School of Life Sciences, École Polytechnique Fédérale de Lausanne, Lausanne CH-1015, Switzerland; School of Chemistry, Raymond and Beverly Sackler Faculty of Exact Sciences, Tel Aviv University, Tel Aviv, 69978 Israel; Protein Production and Structure core facility, School of Life Sciences, École Polytechnique Fédérale de Lausanne, Lausanne CH-1015, Switzerland

## Abstract

Several studies have identified phosphorylation of alpha-synuclein (aSyn) at multiple tyrosine residues (Y39, Y125, Y133, and Y136) within Lewy bodies, which are the pathological hallmarks of Parkinson’s disease (PD). However, understanding the specific role of phosphorylation at each site, or how multiple phosphorylation sites interact, has been challenging. Herein, we present an efficient method that leverages an organometallic Pd-complex to site-specifically attach phosphomimetic groups via Cys-arylation, closely mimicking natural phosphorylation. Using this approach, we successfully incorporated native-like tyrosine phosphorylation mimics into cysteine-containing proteins like synthetic transcription factor Max and recombinant aSyn, in good yields. To demonstrate the method’s versatility, we created a focused library of mono-, di-, and tri-phosphorylated aSyn analogues. This development enabled, for the first time, the investigation of the effect of site-specific phosphorylation at multiple C-terminal tyrosine residues on aSyn fibrillization and aggregation. Our results show that mono-phosphorylation at any of the C-terminal sites has little effect on aggregation, while di- and tri-phosphorylation slows down the process, extending the lag phase compared to wild-type aSyn. Additionally, phosphorylation at Y39 reduces the seeding activity of aSyn fibrils by almost threefold in both mammalian and neuronal models of synuclein pathology. Overall, our strategy provides a rapid, selective, and scalable way to introduce aromatic PTMs, producing homogeneous protein libraries essential for mechanistic studies, biomarker development, and exploring the therapeutic potential of targeting aSyn-phosphorylation for PD and related disorders.

## INTRODUCTION

Post-translational modifications (PTMs) are essential regulatory mechanisms that control a vast array of biological processes crucial for proper cellular function.^1^ The extensive array of these reversible PTMs leads to a significant increase in the structural and functional diversity of the proteome.^1^ It is noteworthy that while different PTMs can modify multiple amino acids, the same amino acid residues often undergo competing PTMs. Of particular interest are tyrosine (Tyr) residues and Tyr-phosphorylation, which play pivotal roles in regulating a variety of processes, including signal transduction, cell growth, differentiation, metabolism, enzymatic activity, and protein-protein interactions.^2^ Aberrant Tyr-phosphorylation has been linked to the onset and progression of various human diseases, including cancer and neurodegenerative disorders.^3^ However, the ability to delineate the precise functional roles of Tyr-phosphorylations in physiological and pathological states remains limited, in part due to the paucity of methods that allow for site-specific introduction of these modifications *in vitro* or within cells.^4^ Despite the noteworthy advancements in total chemical synthesis^5^ and semisynthetic^6^ protein production, implementing late-stage Tyr-phosphorylation remains challenging.

While serine and threonine phosphorylation can be studied using charge-mimicking amino acid substitutions (*e.g.,* serine to aspartate or threonine to glutamate mutations), there are no natural amino acids that replicate the structural and electronic properties of aromatic PTMs (*i.e.,* Tyr-phosphorylation). As a result, site-specific incorporation of these modifications into proteins—particularly in their native, folded states—remains a significant challenge. Enzymatic approaches often suffer from a lack of specificity, leading to heterogeneous protein populations that complicate the analysis of tyrosine modifications and their potential cross-talk with other PTMs. Moreover, our limited understanding of the natural enzymes responsible for writing and erasing tyrosine modifications further impedes insight into their roles in health and disease. Genetic code expansion^7^ offers a potential solution by enabling site-specific incorporation of modified amino acids, but this strategy is technically demanding, particularly when introducing multiple PTMs.^8^ Chemical synthesis and chemoenzymatic ligation have helped overcome some of these barriers and have been instrumental in elucidating PTM functions across various proteins.^9^ However, similar to the semi-synthetic approaches, this is also laborious, time-consuming, and often constrained by the need for denaturing conditions.

To address these limitations, we previously introduced the development of an orthogonal, late-stage protein modification method for the insertion of aromatic PTMs (Tyr-nitration and phosphorylation) into folded proteins via S–C(sp²) bond formation.^10^ Using the neurodegenerative disease-associated protein alpha-Synuclein (aSyn) as a model protein, this strategy enabled the rapid generation of libraries of site-specific and homogeneously nitrated and phosphorylated proteins in desirable yields. However, one limitation of our approach was the insertion of −CH_2_− instead of -O- at the desired Tyr-phosphorylation site, rendering these site-selective mono-phosphorylated aSyn analogues unreactive towards specific antibodies. This made the approach less suitable for tracking them during aggregation inside cells.

In the present study, we significantly advance this methodology by developing Palladium-mediated Cys-arylation chemistry for the site-specific installation of a more “native-like” Tyr-phosphorylation on aSyn. With this method, we first installed the “native-like” Tyr-phosphomimetic at the Y39C position [aSynY39C-P(nat)] on the recombinantly expressed and purified aSyn protein. Our results show that aSyn bearing the native-like Tyr-phosphorylation mimic exhibits similar aggregation properties as native pY39 aSyn prepared using protein semi-synthesis, *i.e.,* significantly reduced aggregation propensity *in vitro* compared to the WT.

Importantly, we used the aSynY39C-P(nat) proteins to determine, for the first time, the effect of bona fide phosphorylation at Y39 on the seeding activity of preformed aSyn fibrils in mammalian cells and neurons and demonstrated that phosphorylated aSyn fibrils at Y39 exhibited less seeding capacity *in vitro,* in WT-aSyn stably-expressing cells, and in primary neurons. Furthermore, we extended this approach to elucidate for the first time the role of site-specific single, double, and triple Tyr-phosphorylation within the C-terminal domain of aSyn. Previous studies using protein semi-synthesis focused on single sites, and attempts to achieve enzyme-mediated site-specific phosphorylation of C-terminal tyrosine residues were unsuccessful. Altogether, the strategy exemplifies a versatile platform that is adaptable to native-like Tyr-phosphorylation of proteins, offering a powerful tool to facilitate the deciphering of protein PTM in health and disease.

## RESULTS AND DISCUSSION

### Facile insertions of phosphotyrosine mimics into proteins through late-stage S-arylation

Selective protein modification provides a versatile approach to diversifying native proteins in ways that are often inaccessible through conventional biological methods.^11^ Palladium (II) oxidative addition complexes (Pd-OACs)^12^ have attracted substantial attention for protein functionalization due to their robust selectivity, high reactivity, and compatibility in aqueous conditions, which enable a broad range of transformations.^13^ For example, we have recently expanded the utility of this approach to incorporate amino acid residues and PTM mimics into peptides and proteins for various applications.^10, 14^ Specifically, we developed Pd-OACs bearing nitro-tyrosine and stable-phosphotyrosine residues, which were successfully transferred into proteins at the desired site(s).^10^ Interestingly, while we showed that the Tyr-nitration mimic was recognized by specific antibodies, the stable-phosphorylation mimic was not recognized by antibodies targeting Tyr-phosphorylation sites. This limits the applicability of the strategy for transferring authentic phosphotyrosine mimics into target protein substrates for biological and cellular studies. Since both PTMs were incorporated via S-arylation chemistry, we hypothesized that the lack of antibody recognition was due to the linker substitution (O–to–CH₂) connecting the phosphate group to the aromatic side chain. To test this hypothesis, we set out to develop a second-generation complex that enables the incorporation of authentic Tyr-phosphorylation into proteins. To this end, we reacted 4-iodophenyl dihydrogen phosphate with (COD)Pd(CH₂SiMe₃)₂ in the presence of the RuPhos ligand to yield the desired product (Pd-OAC-NP) after 4 hours (Figure 1A, Supporting information section 2.2). To assess the reactivity of the isolated complex, we initially tested its potential to transfer the phosphotyrosine mimic into a 9-mer model peptide, P1 (KLYACSRYL), bearing a single Cys residue. Remarkably, treatment of P1 with Pd-OAC-NP (4 equiv) in 20 mM Tris buffer (pH 7.5) containing 5% DMF as a cosolvent led to quantitative conversion to the desired modified product P1-P(nat) within 10 minutes at room temperature, as confirmed by LC−MS analysis (Figure 1B-C, Figure S1). These results demonstrate that the Pd-OAC-NP complex can efficiently install the target phosphotyrosine mimic into unprotected peptides under mild conditions, highlighting its potential for site-specific functionalization of complex macromolecules.

**Figure 1.**
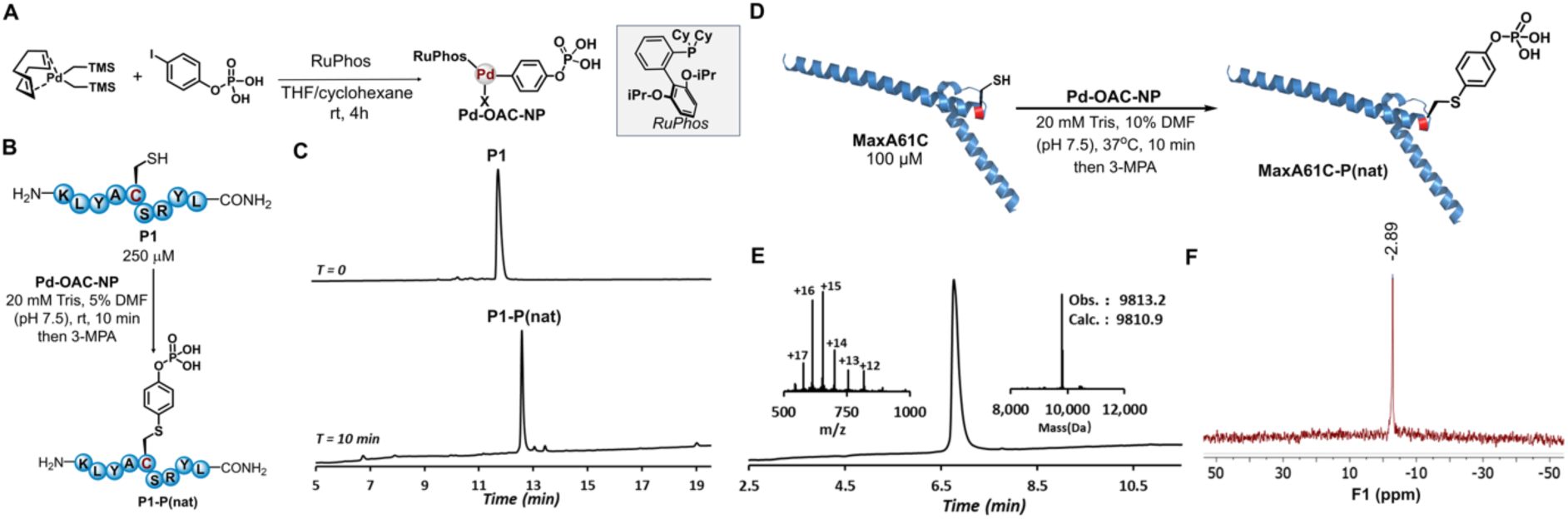
Late-stage insertion of Tyr-phosphorylation mimic to unprotected peptides and proteins using the palladium (II) strategy. (A) Schematic representation of the synthesis of palladium (II) oxidative addition complexes bearing Tyr-phosphorylation mimic (Pd-OAC-NP). (B) Schematic representation of the site-specific installation of Tyr-phosphorylation mimic to the unprotected peptide. (C) Analytical HPLC trace of the crude reactions after 10 min. The reactions were quenched with 3-mercaptopropionic acid (3-MPA) before HPLC analysis. Analytical HPLC was performed by using 0.05% trifluoroacetic acid in water and acetonitrile as the mobile phases. (D) Site-specific installation of Tyr-phosphorylation mimic into MaxA61C protein. (E) LC-MS analysis of isolated phosphorylated product MaxA61C-P(nat). LC-MS was performed using 0.1% formic acid in water and acetonitrile as the mobile phases. Modified protein was obtained in the protonated form. (F) ^31^P NMR spectrum of MaxA61C-P(nat) in D_2_O. X: 4-iodophenyl dihydrogen phosphate.

We next focused on site-selective installation of Tyr phosphorylation in proteins. As a model system, we applied this strategy to modify the DNA-binding domain of the Max transcription factor (TF), engineered with a single Cys residue (MaxA61C). Treatment of MaxA61C with Pd-OAC-NP under our optimized reaction conditions (20 mM Tris buffer, 10% DMF, pH 7.5, 37 °C) resulted in the formation of the desired mono-phosphorylated product, MaxA61C-P(nat), after 10 min, with 56% isolated yield after RP-HPLC purification (see Supporting information, Section 4.1 for details). These results demonstrate the efficient and selective transfer of the Tyr-phosphorylation mimic to the target protein in good yield and excellent purity, as confirmed by LC-MS and ^31^P-NMR (Figure 1D-F, Figure S3 and S27). To evaluate the chemical stability of the installed phosphorylation mark, MaxA61C-P(nat) was incubated in phosphate-buffered saline (PBS, pH ∼7.0) at 37 °C. LC-MS analysis showed no detectable degradation over a 5-day period (see Supporting Information, Section 4.2, Figure S4). Notably, the reaction can be performed on the milligram scale, enabling the preparation of homogeneously modified proteins in sufficient quantities for in-depth structural, biochemical, and biological studies aimed at elucidating the functional relevance of site-specific Tyr-phosphorylation.

### Site-selective insertion of phosphotyrosine mimic into native alpha-Synuclein protein

To further evaluate the efficacy of our phosphomimetic in replicating the effects of authentic Tyr-phosphorylation PTM, we extended our approach to the synaptic protein aSyn.^15^ We used again aSyn as a model system for the following reasons; 1) several native site-specifically phosphorylated forms of the protein have been produced using protein semi-synthesis and the aggregation properties of these proteins have been extensively characterized, thus enabling direct comparison to our native Tyr-phosphorylation mimics; 2) aSyn is among the most extensively targeted proteins for developing diagnostic and therapeutic strategies for the treatment of PD and other neurodegenerative diseases (synucleinopathies);^16^ 3) while effect of phosphorylation at Tyrosine residues on the aggregation of monomeric aSyn have been studied, the effect of phosphorylation on the seeding activity of fibrils, a process that is central to pathology formation and spreading in the brain, has not been assessed thoroughly; 4) no studies have explored the effect of site-specific Tyr-phosphorylation at multiple tyrosine residues, a process that has been observed in PD brain pathology and 5) converging evidence points to PTMs, including phosphorylation residues, as critical regulators of their physiological and pathogenic properties, including aggregation, seeding, and pathology spreading.^17^

### Synthesis of Tyr-phosphorylated aSyn mimics

In our previous study, we observed that the site-specific Tyr →Cys mutation at position 39 of aSyn did not change the aggregation properties of the protein compared to WT-aSyn.^10^ Therefore, in this context, we used aSyn-Y39C mutant as a starting material to produce our “native-like” phospho mimetics of aSyn [Y39C-P(nat)]. The WT-aSyn and aSyn-Y39C proteins were expressed and purified as per our previous report.^10^ Next, we employed our designed Pd(II)OAC described above to produce homogeneously phosphorylated aSyn at Y39 with high fidelity (Figure 2B). Briefly, we initially treated aSyn-Y39C with Pd-OAC-NP using our optimized reaction conditions in 20 mM Tris buffer and 5% DMF (pH 7.5) at 37°C for 15 min. This provided the desired monophosphorylated product aSyn-Y39C-P(nat) in 81% isolated yield after RP-HPLC purification (see the Supporting Information section 18.1). The identity and purity of the phosphorylated mimic were confirmed by LC-MS (Figure 2C, S7) and SDS-PAGE analysis (Supporting Information, Figure S13).

**Figure 2.**
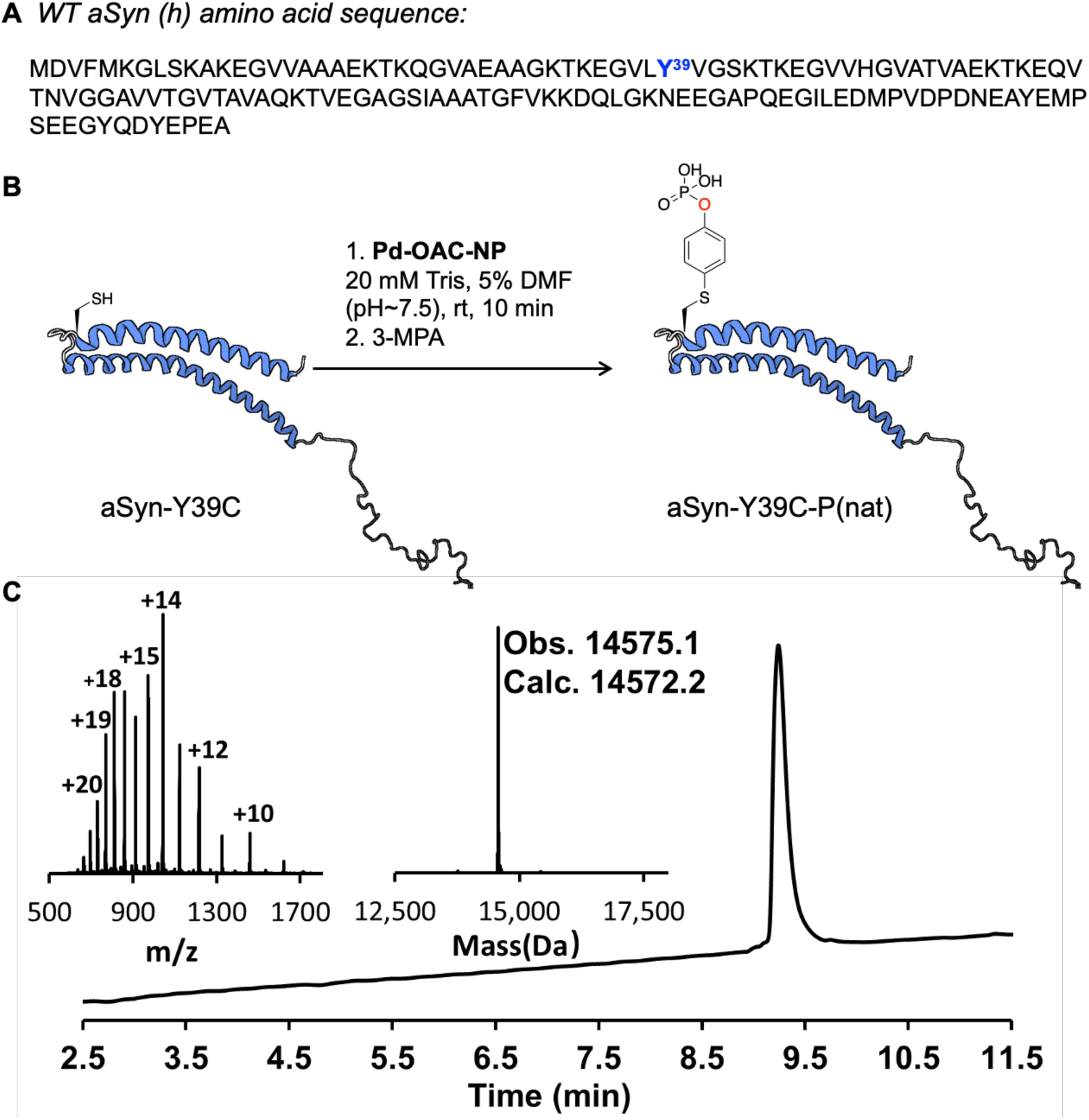
Site-specific phosphorylation of aSyn at Tyr-39. (A) The amino acid sequence of human aSyn wild type (WT-aSyn (h)). The highlighted Tyr^39^ site is site-specifically mutated to a Cys residue and used as the corresponding phosphorylation site. (B) Synthesis of monophosphorylated Y39C-P(nat) analogue from Y39C mutant. (C) LC-MS analysis of the monophosphorylated Y39C-P(nat) protein.

### Phosphorylation at Residue 39 Significantly Inhibits aSyn Aggregation *In Vitro*

Next, we assessed the impact of the “native-like” phospho mimetic at Y39 (for aSyn-Y39C-P(nat)) on the aggregation of aSyn using the established thioflavin T (ThT)-based fluorescence assay.^18^ The kinetics curve clearly showed that the aSyn-Y39C-P(nat) exhibited delayed aggregation (Figure 3B), with a much lower ThT signal compared to WT-aSyn (Figure 3A), even after 100 hours. These results correlated well with the sedimentation assay^19^ (Figure S16A-B), which indicated that nearly 80-90% of the aSyn-Y39C-P(nat) remained soluble (up to 100 h), as assessed by monitoring monomer loss post-aggregation (Figure 3C). Consistent with the ThT and sedimentation studies, monitoring the aggregation by TEM over time also revealed delayed aggregation (Figure 3D), but no significant differences were found in the morphological properties of the aggregates (Figure 3G-H). The TEM image analysis revealed that both the WT and Y39C-P(nat) fibrils are polymorphic in nature with twisted single filaments and stacked multifilament structures, as shown in Figure 3G-H. Using the EM micrographs to quantify and compare their fibril properties showed that there are no significant differences in the length of the two fibrils. Because of the presence of multifilament structures, the average width of the WT fibrils is higher than the average diameter of the Y39C-P(nat) fibrils (Figure 3E-F).

**Figure 3.**
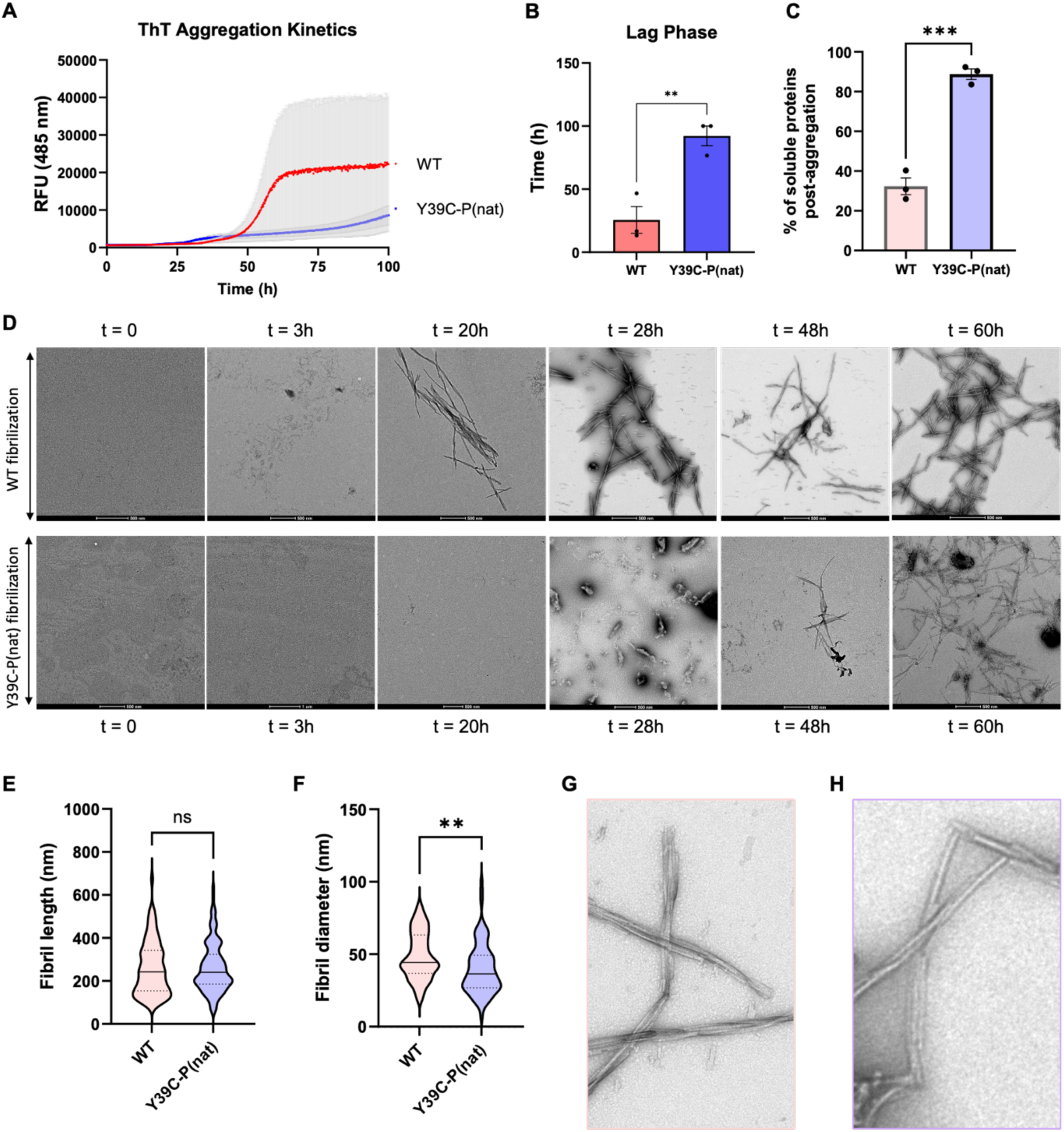
Y39C-P(nat) aggregates much more slowly than WT. (A) Aggregation kinetics of WT and phosphorylated Y39C-P(nat) at 20 µM initial concentration monitored by ThT fluorescence. RFU, relative fluorescence units, n = 2. (B) Bar graph of lag phase extracted from the ThT aggregation kinetics shown by (A) (mean ± SEM, n = 3). (C) Solubility assay showing the percentage of soluble protein post-aggregation of all the aSyn analogues (at the final time point). (D) Electron micrographs of time-dependent WT and Y39C-P(nat) aggregation from 20 µM. Scale bars, 500 nm. (E) Comparison of the WT and Y39C-P(nat) fibrils with the average WT fibril length ∼259 nm and average WT fibril length ∼262 nm. (F) Comparison of the WT and Y39C-P(nat) fibrils, with the average WT fibril width ∼49 nm and the average Y39C-P(nat) fibril width ∼40 nm. (G, H) Zoomed EM micrographs of WT and Y39C-P(nat) fibrils showing twists and lateral stacking.

The decreased aggregation tendency for Y39C-P(nat) aligns with our earlier findings using semi-synthetically prepared pY39-aSyn, which validates our native-like phosphomimetic approach.^20^ Next, we sought to determine if Y39C-P(nat) proteins could be detected using antibodies that are specific to the bona fide pY39-aSyn. This is important to determine if these proteins could be used and selectively monitored in cell culture or *in vivo* studies. We compared the “native-like” [Y39C-P(nat)] and pY39 proteins using dot blot analysis. As shown in supporting Figure S14Ai, both pY39 and Y39C-P(nat) are recognized by pY39-aSyn antibodies. Moreover, when the Y39C-P(nat) analogue is dephosphorylated *in vitro* in the presence of phosphatase (see the Supporting Information Figure S14Bi), the protein is no longer detected by the pY39 antibodies. Together, these observations support that the Y39C-P(nat) construct can be used in cellular assays as an alternative to semi-synthetically prepared or mutated, phosphomimicking amino acid residues (*e.g.,* Y39E or Y39D), which are not accurate mimics of tyrosine phosphorylation.

### Generation of PFFs and *in vitro* Cross-seeding assay

Studies from our lab and others have shown that PTMs at the level of monomeric or fibrillar aSyn can dramatically influence the aggregation, seeding, and propagation of aSyn.^17^ In particular, we recently showed that O-linked-N-Acetylglucosaminylation (O-GlcNAcylation) at the Ser87 position^17b^ or nitration of all tyrosine residues on aSyn^21^ diminishes the seeding activity of aSyn *in vitro*, in neurons, and *in vivo*. Previous attempts to investigate the effect of site-specific phosphorylation at Y39 on aSyn seeding activity have relied mainly on the use of phosphomimicking mutations (Y → E)^22^ or mutating all C-terminal tyrosine residues (Y → F) to allow for site-specific phosphorylation at Y39.^23^ Consequently, we proceeded to evaluate the *in vitro* seeding activity of fibrils generated through the *in vitro* fibrillization of Y39C-P(nat). We generated the preformed fibrils (PFFs) from WT, Y39C mutant, and Y39C-P(nat), Figure S18.^19^ The fibrils were sonicated to achieve a median fibril length of ∼50–100 nm with an average width of ∼15 nm (Figure 4A-B) and were characterized by ThT and TEM. To ensure an equal amount of PFF seeds was used for all three cases in the seeding experiments, we measured the effective concentration of the PFFs by determining the ratio of soluble to fibrillar aSyn species using ultracentrifugation-based sedimentation and filtration assays. The separated PFFs were resuspended and subjected to concentration measurement via a standard BCA assay.

**Figure 4.**
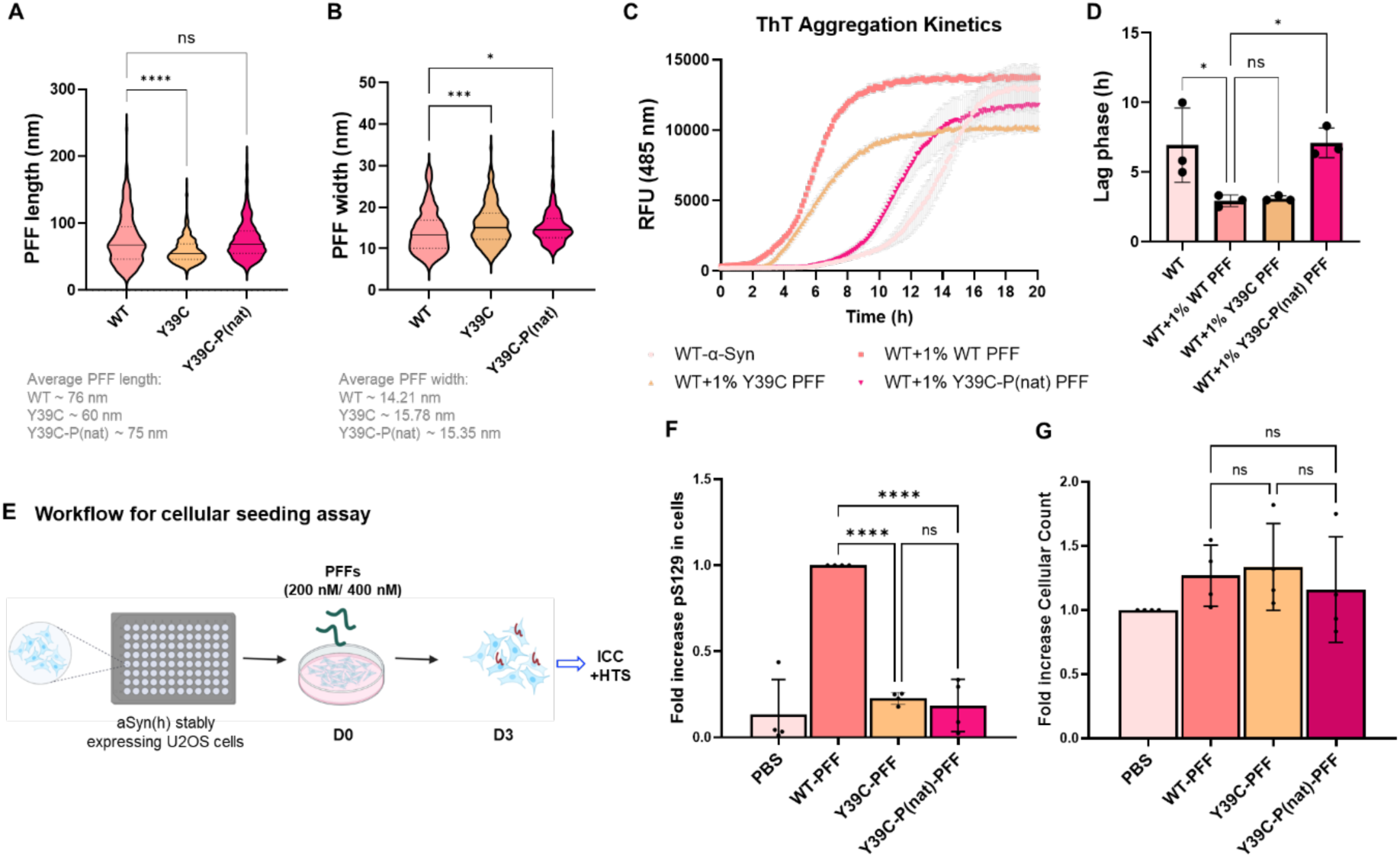
(A) Quantification of the length of aSyn WT, Y39C, and Y39C-P(nat) PFFs. (B) Quantification of the width of aSyn WT, Y39C, and Y39C-P(nat) PFFs. (C) *In vitro* cross-seeding assay via ThT fluorescence-based aggregation kinetics plot. (D) Bar graph of lag phase from the aggregation kinetics (n = 3, error bars are standard error of the mean). (E) Workflow for the cellular seeding assay to assess the level of pS129 pathology in U2OS cells treated with three types of PFFs. Cells are treated with 200 nM of PFFs separately. Three days post-treatment (D3), the cells were fixed in 4% paraformaldehyde (PFA), and immunocytochemistry (ICC) was performed. (F) Fold Increase of total pS129 level in cells (81a), calculated from cells treated with recombinant human aSyn. (one-way ANOVA, N = 4). (G) Fold Increase of cellular count for each condition (Phalloidin), calculated from cells treated with PBS. (one-way ANOVA, N = 4).

As expected, and shown in Figure 4C, seeding with WT PFFs significantly increased the aggregation rate of the WT-monomers compared to the non-seeded WT, as evidenced by the shift of the lag phase from ∼7 h to ∼3 h. In contrast, seeding with Y39C-P(nat) fibrils only modestly modified the lag phase of WT aggregation (∼8 h). WT and Y39C PFFs exhibited similar lag phase and seeding activity, further confirming that the presence of the Cys residue does not influence aSyn aggregation of fibril seeding activity (Figure 4C). Altogether, our findings establish that phosphorylation at Y39 diminishes the seeding activity of aSyn fibrils *in vitro*.

### Y39C-P(nat) PFFs exhibited significantly less seeding activity than WT PFFs in U2OS cells and neurone-based cellular models

To validate our *in vitro* results, we next sought to assess the seeding activity of Y39C-P(nat) in cellular models of aSyn seeding and pathology formation. We established a cellular seeding model with stably expressed WT-aSyn (h) U2OS cells (Fig. S19) to study the effects of PFFs in a more time-efficient manner (Figure 4E). The cells were treated with 200 nM of three distinct aSyn-PFF species— WT, Y39C, and Y39C-P(nat) — or with PBS as a negative control. The extent of seeding activity was assessed by monitoring and quantifying the level of phosphorylated aSyn aggregates using pS129-specific antibodies. Immunocytochemistry (ICC) followed by fluorescence imaging confirmed the formation of pS129-positive aggregates after 72h of PFF treatment. Quantification of the pS129 levels by high-throughput screening (HTS) and high content analysis (HCA) demonstrated that lower levels of pS129-seeded aggregates were formed in cells treated with Y39C-P(nat) PFFs than in those treated with WT PFFs (Figure 4F). Our quantitative data further indicated that the decrease in aSyn aggregate count is not due to cell death/lower cell count in the Y39C-P(nat) PFF-treated wells, as cellular counts were comparable across all conditions tested (Figure 4G). Notably, the Y39C mutant PFF also resulted in a marked decrease in pS129 pathology, which is similar to Y39C-P(nat) PFFs, that is approximately one-fourth of the seeding capacity of WT PFFs. Interestingly, in the *in vitro* seeding assay, the Y39C mutant PFFs did not exhibit such a pronounced effect on soluble WT-aSyn.

Subsequently, we conducted similar seeding studies in primary hippocampal neurons in the absence of aSyn overexpression. In this model, low nanomolar concentrations of PFFs (70 nM) added to primary neuronal cultures are sufficient to induce the formation of intracellular fibrils that bear similar biochemical properties (PTM signatures) as fibrils found in the brains of PD patients.^17b–c, 22b^ Importantly, the newly seeded fibrils in this model undergo remodelling, further modifications, and transition to LB-like inclusions at later stages, which resemble LBs in PD brains at both biochemical and ultrastructural levels.^17c, 24^ The seeding capacities were compared in wild-type (WT) murine hippocampal neurons by administering the respective PFFs to WT primary cultures on day 7 (DIV7). Their seeding ability to initiate the formation of pS129 pathology was assessed at day 14 (D14) post-treatment (Figure 5A). Quantification of pS129 levels via HTS and followed by HCA revealed that a greater number of pS129-seeded aggregates were formed in hippocampal neurons at D14 that correspond to WT PFFs compared to those treated with Y39C-P(nat) PFFs (Figure 5A-B). Also, in this case, the significant drop in aggregate level is not due to the low cell count in the Y39C-P(nat) PFF-treated regions, as the cellular count remains identical for each treatment/condition tested, including PBS control (Figure 5C).

**Figure 5.**
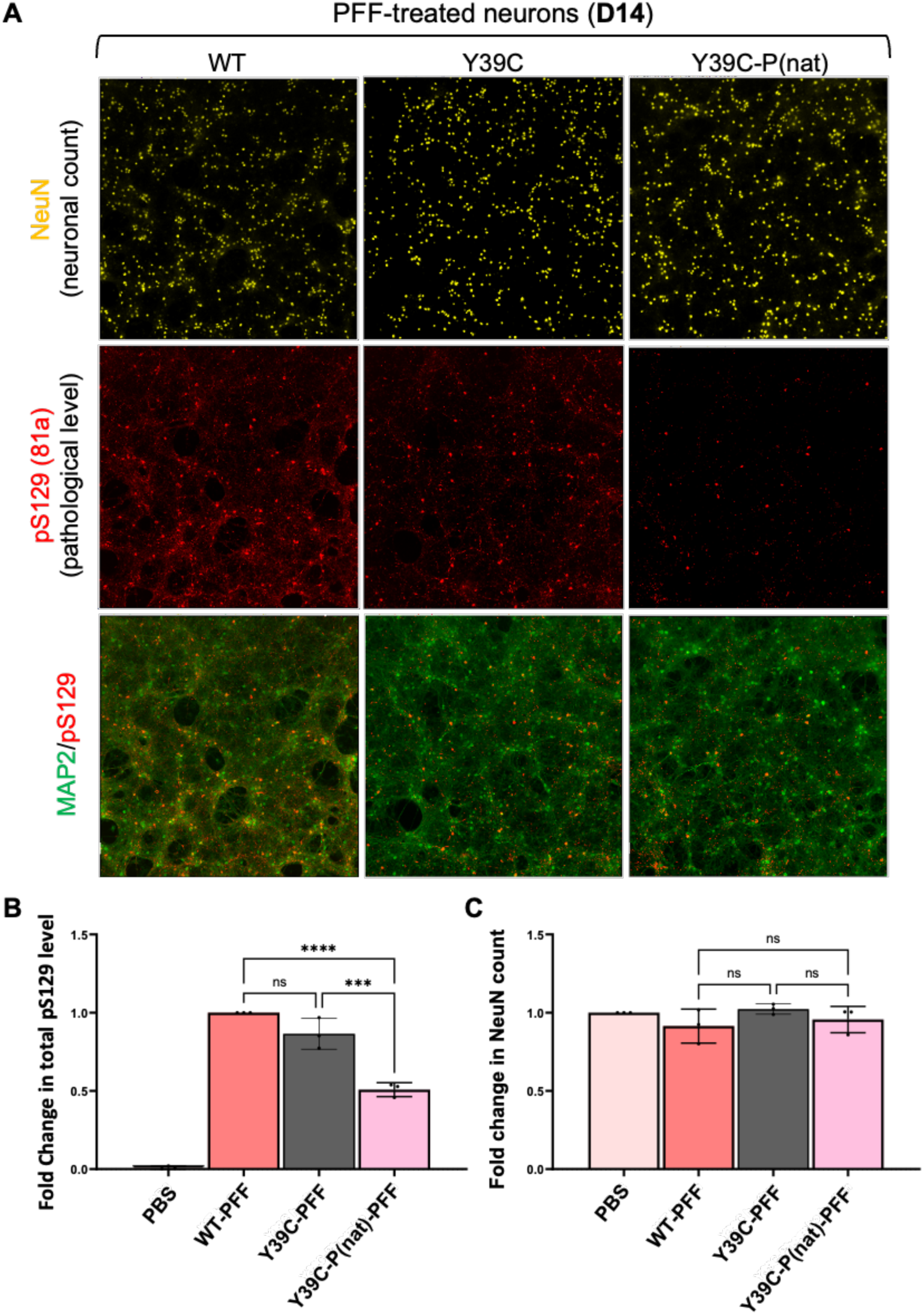
Level of pS129 pathology in WT neurons treated with PFFs on DIV7. WT neurons are treated with 70nM of the respective PFFs. Fourteen days post-treatment (D14), the neurons were preserved in 4% PFA, and ICC was subsequently conducted. (A) Representative images of the D14 PFF-treated neurons. (B) The fold increase of total phosphorylated serine 129 (pS129) level in cells (81a) is assessed, derived from neurons treated with recombinant human alpha-synuclein. A one-way ANOVA was conducted with N = 3. (C) The fold increase of cellular count for each condition (MAP2) is calculated based on neurons treated with PBS. A one-way ANOVA was performed, with N = 3.

Interestingly, unlike what we observed in the U2OS cells, fibrils derived from WT and Y39C mutant aSyn showed similar seeding activity in neurons, consistent with the *in vitro* seeding studies. Together, these observations establish that phosphorylation at Y39 inhibits the seeding activity of aSyn fibrils and suggest that the neuronal seeding model, where seeding is assessed in the absence of aSyn overexpression, may be a more reliable approach to studying the effects of the PTM(s) on aSyn seeding and pathology formation.

### Phosphorylation and hyperphosphorylation at the C-terminal domain of aSyn

To determine if our approach could be extended to other Tyr residues within aSyn or to investigate the effect of Tyr-phosphorylation at multiple residues, we next assessed the effect of introducing native-like phosphomimetics site-specifically at the C-terminal Tyr residues of aSyn. Phosphorylated aSyn at Tyr residues (Y125, Y133, or Y136) have been detected in Lewy bodies (LBs) and glial cytoplasmic inclusions (GCIs).^23a, 25^ Furthermore, recent studies from our group have also validated the presence of hyperphosphorylated (mainly pY133 and pY136) aSyn aggregates in the neuronal seeding model and in the PFF-injected *in vivo* model.^25b^ However, the effect of site-specific Tyr-phosphorylation on aSyn aggregation has only been assessed at Y125 using semisynthetic pY125 proteins.^26^ This is primarily due to the lack of facile approaches that enable site-specific modification of Tyr residues or the introduction of multiple phosphorylated Tyr residues. Several kinases have been reported to phosphorylate aSyn Tyr sites *in vitro* and via cellular studies.^27^ For example, the tyrosine kinase c-Abl phosphorylates aSyn, primarily at Y39 and partially at Y125 *in vitro*, in neuroblastoma cell lines and primary cultures of mouse cortical neurons.^26a^ Tyrosine-phosphorylation at Y125 has been reported by Src, Fyn, Syk, Lyn, and c-Frg tyrosine kinases.^28^ Syk kinases are also found to phosphorylate Y133 and Y136 positions of aSyn.^28c^ However, no kinases have been shown to selectively phosphorylate aSyn at single or multiple, namely double, C-terminal tyrosine residues. Although this is possible using protein synthesis and semisynthetic approaches,^29^ the introduction of native-like Tyr-phosphomimetics into recombinant proteins enables rapid generation and assessment of a large number of modifications and the crosstalk between multiple PTMs.

Towards this aim, we expressed and purified the respective aSyn Cys mutants, *i.e.,* Y125C/Y133C (Figure S5), and Y125C/Y133C/Y136C (Figure S6) and reacted them with Pd-OAC-NP, utilizing the previously optimized reaction conditions to generate homogeneously di- and tri-phosphorylated analogues Y125C/Y133C-P(nat) and Y125C/Y133C/Y136C-P(nat) in 45% and 54% isolated yields respectively (see Figures 6B and 6C). To enable comparison of the aggregation of the multi-phosphorylated analogues with the corresponding mono-phosphomimetic aSyn construct(s), we also synthesized aSyn-Y125C-P(nat), aSyn-Y133C-P(nat), and aSyn-Y136C-P(nat) in 89%, 87%, and 82% isolated yields from the respective Cys-mutants aSyn (see the Supporting Information for the synthesis of mono- and multi-phosphorylated analogues, Section 18, Figure S8-S12).

**Figure 6.**
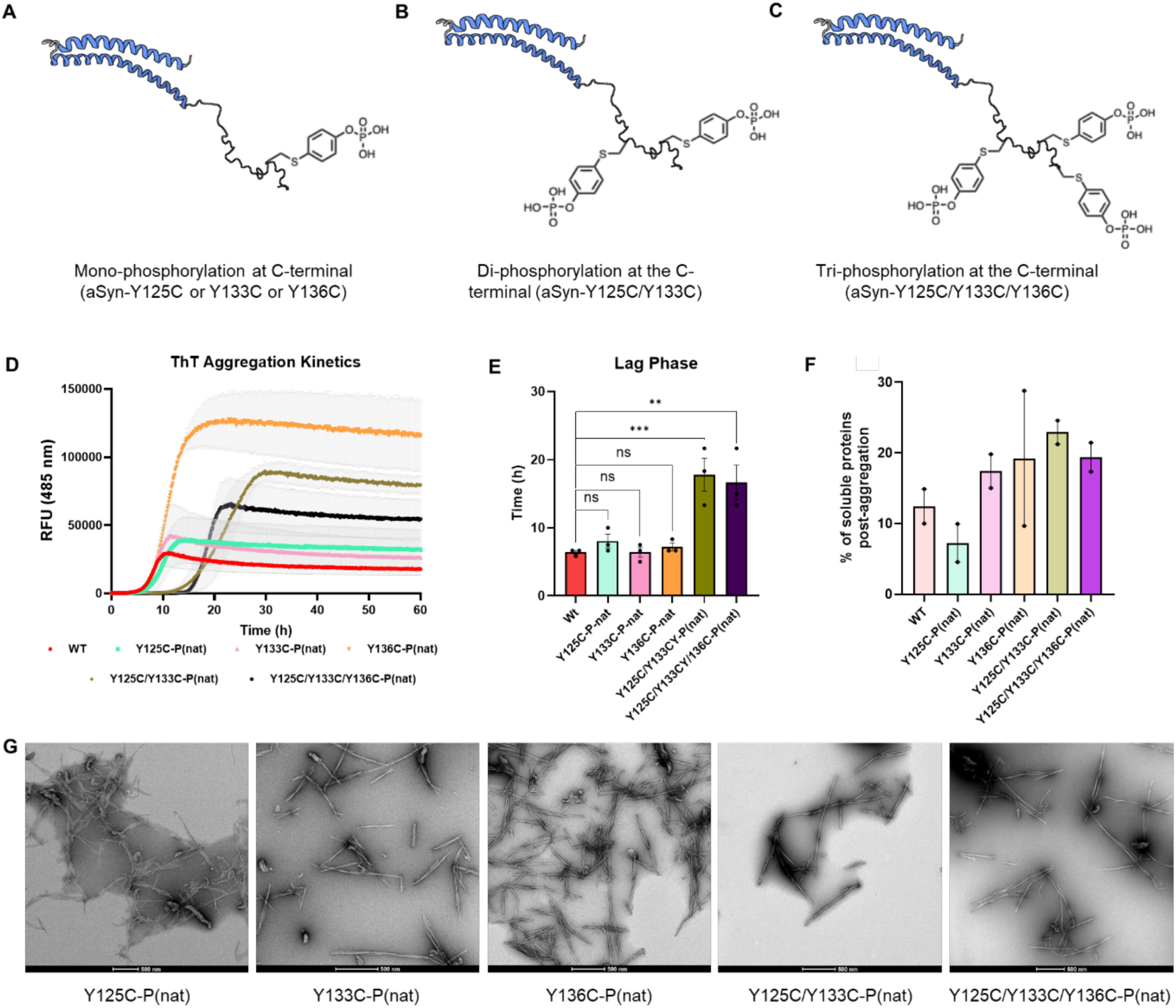
aSyn C-terminal mono- and hyper-phosphorylation. (A) Synthesis of Y125C-P(nat) or Y133C-P(nat) or Y136C-P(nat). (B) Y125C/Y133C-P(nat) and (C) Y125C/Y133C/Y136C-P(nat) analogues. (D) Comparison of aggregation kinetics of WT, mono-, and hyperphosphorylated aSyn analogues at an initial concentration of 20 µM, monitored by ThT fluorescence. RFU, relative fluorescence units (mean ± SEM, n = 2). (E) Bar graph of lag phase extracted from the ThT aggregation kinetics (mean ± SEM, n = 3). (F) Bar graph of quantified soluble protein percentage post-aggregation from sedimentation assay following ThT aggregation assay (mean ± SEM, n = 2). (G) Electron micrographs of the mono-phosphorylated, di-phosphorylated, and tri-phosphorylated aSyn analogues, respectively (scale bars, 500 nm for all the TEM images). See supporting information for zoomed images.

To evaluate the impact of mono vs multi-phosphorylation on aSyn aggregation, we conducted a comparative analysis of the fibrillization kinetics of WT-aSyn, mono-phosphomimetic analogues [Y125C-P(nat), Y133C-P(nat), Y136C-P(nat)], and multi-phosphomimetic analogues [Y125C/Y133C-P(nat), Y125C/Y133C/Y136C-P(nat)] via the established ThT assay (Figure 6D). As shown in Figure 6D, the C-terminally mono-phosphomimetic analogues [Y125C-P(nat), Y133C-P(nat), Y136C-P(nat)] behaved similarly to the WT-aSyn, consistent with our previous findings using semi-synthetically prepared pY125.^26^ This again validates our approach and the utility of using “native-like” phosphomimetic group(s) to investigate the role of Tyr-phosphorylation in regulating aSyn aggregation. On the other hand, di- and tri-phosphomimic analogues [Y125C/Y133C-P(nat), Y125C/133C/Y136C-P(nat)] exhibited relatively slower aggregation with a significantly longer lag phase compared to the WT-aSyn (Figure 6D and 6E). The aggregation profile also matched with the respective sedimentation assay (Figure S16C-H). To analyze the properties of the fibrils, we quantified the length and diameter of each fibril. It revealed that the C-terminal mono-, di-, and tri-phosphomimetic analogues are significantly longer than the WT fibrils (see Figure S17A for details). Interestingly, the width of the di- and tri-phosphomimetics is considerably smaller than that of the WT or the C-terminally mono-phosphomimetic analogues (Figure S17B). Similar to the WT fibril, the C-terminally mono-phosphomimetic analogues possess stacked multifilaments (Figure S17C-E, zoomed EM images); however, the multiphosphomimetic analogues mainly exhibited long and single filaments (Figure S17F-G, zoomed EM images) with an average length of 495 nm and 467 nm and an average diameter of 31 nm and 20 nm, for di- and tri-phosphomimetic fibrils, respectively. One probable reason for such observation could be that the presence of multi-Tyr-Phosphorylations inhibits the lateral overlapping/stacking of the fibrils.

It is worth noting that the unstructured C-terminal tail region of aSyn is fully solvent-exposed, contains multiple Tyr residues, and is highly enriched in acidic residues. Consequently, this could explain why substituting a single Tyr residue with the phosphomimetic unit did not substantially alter the aggregation profile compared to WT-aSyn. Nevertheless, introducing the di- and tri-phosphorylation also delayed the aggregation of aSyn, which is of particular interest. These results underscore the critical importance of conducting further studies to investigate the crosstalk between phosphorylation at different tyrosine residues and the mechanisms by which hyperphosphorylation influences aSyn aggregation and seeding activity. The approach outlined and validated here paves the way for achieving this goal by facilitating the rapid generation of libraries of site-specific Tyr-phosphorylated aSyn at single or multiple residues.

## CONCLUSION

The site-specific incorporation of aromatic PTMs into recombinant proteins offers a powerful approach to generate homogeneous protein analogs bearing novel aromatic PTMs, such as Tyr phosphorylation. In this work, we developed a method for chemical modification of recombinant proteins, leveraging an organometallic Pd-complex to site-specifically install phosphomimetic groups as Cys-arylation that closely resemble natural phosphorylation. Using this method, we successfully incorporated Tyr-phosphorylation mimics into several peptides and protein targets, including the transcription factor Max and the synaptic protein aSyn, with high yields and purities on a multimilligram scale. With this approach, we generated native-like Y39-phosphorylated aSyn, validated the analogue biochemically, and thoroughly examined the effect of aSyn Y39-phosphorylation in aggregation, *in vitro* seeding, and pathology spreading in cellular disease models. To the best of our knowledge, this is the first report of such a thorough analysis, and we find that this PTM mark substantially reduces aSyn seeding in neurons. Finally, this approach enabled site-specific investigation of C-terminal phosphorylation at multiple tyrosine residues, providing new insights into how differential phosphorylation at these residues modulates aSyn fibrillization kinetics and aggregate morphology. Previous studies on the effect of phosphorylation at Y39 have shown that it dramatically slows the aggregation of monomeric aSyn *in vitro.*^10^ These effects on the aggregation of monomeric aSyn are consistent with previous studies from our lab, site-specifically phosphorylated aSyn at Y39 produced via protein semisynthetic approaches.^20^

Previous report from Brahmachari *et al.* showed that co-expression of kinase-active c-Abl and WT-aSyn in HEK293T cells enhanced aSyn aggregation—a phenotype suppressed by kinase-dead c-Abl or a phospho-deficient Y39F mutant, implying the Y39 phosphorylation promotes aggregation and pathology^23b^. However, it is worth noting that in this work, the extent of Y39 phosphorylation was not quantitatively assessed—the emergence of puncta upon 48h of transfection was marked as aggregates. Therefore, our method, which utilized established disease models to check seeding properties, is not comparable to the work of Brahmachari *et al.* Furthermore, we previously showed that C-Abl phosphorylates both Y39 and Y125 on aSyn, which can also affect the result and interpretation.^23a^ Another work by Zhao *et al.*^30^ reported that pY39 enhances pathology in rat primary cortical neurons; however, their system/model is limited due to a lack of proper WT-aSyn seeding phenomena captured in the time frame monitored, which prevents a direct comparison with our approach using hippocampal mouse neurons as aSyn seeding model.

It is worth noting that the unstructured C-terminal tail region of aSyn is fully solvent-exposed, contains multiple Tyr residues, and is highly enriched in acidic residues. Consequently, this could explain why substituting a single Tyr residue with the phosphomimetic unit did not substantially alter the aggregation profile compared to WT-aSyn. Nevertheless, introducing di-and tri-phosphorylation also delayed aSyn aggregation, which is of particular interest. Although previous studies showed that aSyn inclusions can be stained with antibodies against different tyrosine phosphorylated C-terminal residues,^25b^ it remains unclear whether these PTMs occur on the same molecules. These results underscore the critical importance of conducting further studies to map aSyn C-terminal PTM at a single molecule level and investigate the crosstalk between phosphorylation at different tyrosine residues and the mechanisms by which hyperphosphorylation influences aSyn aggregation and seeding activity. The approach outlined and validated here paves the way to achieving this goal by facilitating the rapid generation of libraries of site-specific Tyr-phosphorylated aSyn at single or multiple residues.

As our methodology provided access to site-specific mono- and multi-phosphorylated aSyn variants, enabling the systematic dissection of the effects of individual and multiple tyrosine modifications and PTMs crosstalk. This robust, scalable chemical strategy overcomes key challenges in protein modification and opens the door to broader mechanistic studies, biomarker development, and the exploration of the therapeutic potential of targeting aSyn phosphorylation for the treatment of PD and other neurodegenerative diseases.

## Supporting information

The supporting information includes synthesis and methods, as well as biochemical, biophysical and biological characterization of all compounds.

## ASSOCIATED CONTENT

### Supporting Information

#### Author Contributions

The manuscript was written through the contributions of all authors. All authors have given approval to the final version of the manuscript. ‡These authors contributed equally.

#### Notes

The authors declare no competing financial interest.

## ACKNOWLEDGMENT

This study was supported by the EPFL (H.A.L., S.M., A.V.G., A.L.M., and Y.J.), grant 1398 (S.M.), and ISF Grant 493/23, the Neubauer Foundation, and the Council for Higher Education (M.J.). We thank the Protein Production and Structure Core Facility for the expression and purification of the aSyn WT and mutant proteins used in this study (B.B.) and the Interdisciplinary Centre for Electron Microscopy for providing access to the electron microscopes.

## Table of Contents graphic

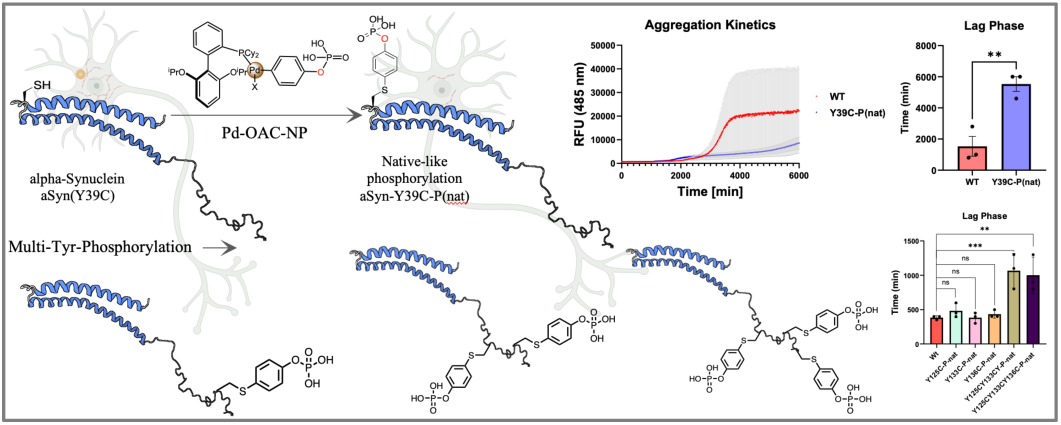

## REFERENCES

1. (a) Barber, K. W.; Rinehart, J. The ABCs of PTMs. Nat. Chem. Biol. 2018, 14 (3), 188−192. (b) Walsh, C. T.; Garneau-Tsodikova, S.; Gatto, G. J., Jr. Protein posttranslational modifications: the chemistry of proteome diversifications. Angew. Chem., Int. Ed. 2005, 44 (45), 7342−7372.

2 Hunter, T. The Genesis of Tyrosine Phosphorylation. Cold Spring Harbor Perspect. Biol. 2014, 6 (5), No. a020644.

3 Zhong, Q.; Xiao, X.; Qiu, Y.; Xu, Z.; Chen, C.; Chong, B.; Zhao, X.; Hai, S.; Li, S.; An, Z.;, et al. Protein posttranslational modifications in health and diseases: Functions, regulatory mechanisms, and therapeutic implications. Medcomm. 2023, 4 (3), No. e261.

4 (a) Spicer, C. D.; Davis, B. G. Selective chemical protein modification. Nat. Commun. 2014, 5, 4740. (b) Krall, N.; da Cruz, F. P.; Boutureira, O.; Bernardes, G. J. Site-selective protein-modification chemistry for basic biology and drug development. Nat. Chem. 2016, 8 (2), 103−113.

5 Bondalapati, S.; Jbara, M.; Brik, A. Expanding the chemical toolbox for the synthesis of large and uniquely modified proteins. Nat. Chem. 2016, 8 (5), 407−418.

6 (a) Holt, M.; Muir, T. Application of the protein semisynthesis strategy to the generation of modified chromatin. Annu. Rev. Biochem. 2015, 84, 265−290. (b) Wang, Z. A.; Cole, P. A. Methods and Applications of Expressed Protein Ligation. Methods Mol. Biol. 2020, 2133, 1−13.

7 (a) Chin, J. W. Expanding and reprogramming the genetic code. Nature 2017, 550 (7674), 53−60. (b) Hoppmann, C.; Wong, A.; Yang, B.; Li, S. W.; Hunter, T.; Shokat, K. M.; Wang, L. Site-specific incorporation of phosphotyrosine using an expanded genetic code. Nat. Chem. Biol. 2017, 13 (8), 842.

8 (a) Lang, K.; Chin, J. W. Cellular incorporation of unnatural amino acids and bioorthogonal labeling of proteins. Chem. Rev. 2014, 114 (9), 4764−4806. (b) Xie, J. M.; Schultz, P. G. Innovation: A chemical toolkit for proteins - an expanded genetic code. Nat Rev Mol Cell Bio. 2006, 7 (10), 775−782.

9. (a) Kent, S. Chemical protein synthesis: Inventing synthetic methods to decipher how proteins work. Bioorg. Med. Chem. 2017, 25 (18), 4926−4937. (b) Hackenberger, C. P.; Schwarzer, D. Chemo-selective ligation and modification strategies for peptides and proteins. Angew. Chem., Int. Ed. 2008, 47 (52), 10030−10074. (c) Thompson, R. E.; Muir, T. W. Chemoenzymatic Semisynthesis of Proteins. Chem. Rev. 2020, 120, 6, 3051–3126.

10 Lin, X; Mandal, S.; Nithun, R. V.; Kolla, R.; Bouri, B.; Lashuel, H. A.; Jbara, M. A Versatile Method for Site-Specific Chemical Installation of Aromatic Posttranslational Modification Analogs into Proteins. J. Am. Chem.Soc. 2024, 146, 37, 25788–25798.

11. (a) Harel, O.; Jbara, M. Posttranslational Chemical Mutagenesis Methods to Insert Posttranslational Modifications into Recombinant Proteins. Molecules 2022, 27, 4389. 10.3390/molecules27144389. (b) Tilden, J. A. R.; Doud, E. A.; Montgomery, H. R.; Maynard, H. D.; Spokoyny, A. M. Organometallic Chemistry Tools for Building Biologically Relevant Nanoscale Systems. J. Am. Chem. Soc. 2024, 146, 44, 29989–30003. (c) Harel, O.; Jbara, M. Chemical Synthesis of Bioactive Proteins. Angew. Chem. Int. Ed. 2023, 62, e202217716.

12 (a) Vinogradova, E., Zhang, C., Spokoyny, A., et al. Organometallic palladium reagents for cysteine bioconjugation. Nature 2015, 526, 687–691. 10.1038/nature15739. (b) Montgomery, H. R.; Spokoyny, A. M.; Maynard, Heather D. Organometallic Oxidative Addition Complexes for *S*-Arylation of Free Cysteines. Bioconjugate Chem. 2024, 35 (7), 883–889.

13. (a) Jbara, M.; Pomplun, S.; Schissel, C. K. Hawken, S. W.; Boija, A.; Klein, I.; Rodriguez, J.; Buchwald, S. L.; Pentelute, B. L. Engineering Bioactive Dimeric Transcription Factor Analogs via Palladium Rebound Reagents. J. Am. Chem. Soc. 2021, 143, 30, 11788–11798. (b) Jbara, M.; Rodriguez, J.; Dhanjee, H. H.; Loas, A.; Buchwald, S. L.; Pentelute, B. L. Oligonucleotide Bioconjugation with Bifunctional Palladium Reagents. Angew. Chem. Int. Ed. 2021, 60, 12109–12115. (c) Nadal-Bufi, F.; Nithun, R. V.; Moliner, F.; Lin, X.; Habiballah, S.; Jbara, M.; Vendrell, M. Late-Stage Minimal Labeling of Peptides and Proteins for Real-Time Imaging of Cellular Trafficking. ACS Cent. Sci. 2025 11 (1), 66–75. (d) Jbara, M. Transition metal catalyzed site-selective cysteine diversification of proteins. Pure and Applied Chem., 2021, 93 (2), 169–186.

14. (a) Lin, X.; Nithun, R. V.; Samanta, R.; Harel, O.; Jbara, M. Enabling Peptide Ligation at Aromatic Junction Mimics via Native Chemical Ligation and Palladium-Mediated S-Arylation. Org. Lett. 2023, 25, 25, 4715–4719. (b) Lin, X.; Harel, O.; Jbara, M. Chemical Engineering of Artificial Transcription Factors by Orthogonal Palladium(II)-Mediated S-Arylation Reactions. Angew. Chem. Int. Ed. 2024, 63(5), e202317511.

15 (a) Burré, J. The Synaptic Function of α-Synuclein. J Parkinson Dis. 2015, 5 (4), 699−713. (b) Iwai, A.; Masliah, E.; Yoshimoto, M.; Ge, N. F.; Flanagan, L.; Desilva, H. A. R.; Kittel, A.; Saitoh, T. The Precursor Protein of Non-Aβ Component of Alzheimers Disease Amyloid Is a Presynaptic Protein of the Central Nervous System. Neuron 1995, 14 (2), 467−475.

16 (a) Spillantini, M. G.; Goedert, M. The α-synucleinopathies: Parkinson’s disease, dementia with Lewy bodies, and multiple system atrophy. Ann Ny Acad Sci 2000, 920, 16−27. (b) Magalhaes, P.; Lashuel, H. A. Opportunities and challenges of alpha-synuclein as a potential biomarker for Parkinson’s disease and other synucleinopathies. NPJ Parkinsons Dis. 2022, 8 (1), 93.

17 (a) Levine, P. M.; Galesic, A.; Balana, A. T.; Mahul-Mellier, A. L.; Navarro, M. X.; De Leon, C. A.; Lashuel, H. A.; Pratt, M. R. alphaSynuclein O-GlcNAcylation alters aggregation and toxicity, revealing certain residues as potential inhibitors of Parkinson’s disease. Proc Natl Acad Sci U S A 2019, 116 (5), 1511−1519. (b) Balana, A. T.; Mahul-Mellier, A. L.; Nguyen, B. A.; Horvath, M.; Javed, A.; Hard, E. R.; Jasiqi, Y.; Singh, P.; Afrin, S.; Pedretti, R.; et al. O-GlcNAc forces an alpha-synuclein amyloid strain with notably diminished seeding and pathology. Nat Chem Biol 2024, 20 (5), 646−655. (c) Mahul Mellier, A. L.; Burtscher, J.; Maharjan, N.; Weerens, L.; Croisier, M.; Kuttler, F.; Leleu, M.; Knott, G. W.; Lashuel, H. A. The process of Lewy body formation, rather than simply alpha-synuclein fibrillization, is one of the major drivers of neurodegeneration. Proc Natl Acad Sci U S A 2020, 117 (9), 4971−4982. (d) Pan, B.; Shimogawa, M.; Zhao, J.; Rhoades, E.; Kashina, A.; Petersson, E. J. Cysteine-Based Mimic of Arginylation Reproduces Neuroprotective Effects of the Authentic Post-Translational Modification on alpha-Synuclein. J. Am. Chem. Soc. 2022, 144 (17), 7911−7918. (e) Pan, B.; Petersson, E. J. A PARP-1 Feed-Forward Mechanism To Accelerate alpha-Synuclein Toxicity in Parkinson’s Disease. Biochemistry 2019, 58 (7), 859−860. (f) Pan, B.; Rhoades, E.; Petersson, E. J. Chemoenzymatic Semisynthesis of Phosphorylated alpha-Synuclein Enables Identification of a Bidirectional Effect on Fibril Formation. ACS Chem. Biol. 2020, 15 (3), 640−645. (g) Hu, J., Xia, W., Zeng, S. et al. Phosphorylation and O-GlcNAcylation at the same α-synuclein site generate distinct fibril structures. Nat Commun, 2024, 15, 2677. 10.1038/s41467-024-46898-1. (h) Zhang, S., Zhu, R., Pan, B. et al. Post-translational modifications of soluble α-synuclein regulate the amplification of pathological α-synuclein. Nat Neurosci, 2023, 26, 213–225. 10.1038/s41593-022-01239-7.

18 Naiki, H.; Higuchi, K.; Hosokawa, M.; Takeda, T. Fluorometric determination of amyloid fibrils in vitro using the fluorescent dye, thioflavin T1. Anal. Biochem. 1989, 177 (2), 244−249.

19 Kumar, S. T.; Donzelli, S.; Chiki, A.; Syed, M. M. K.; Lashuel, H. A. A simple, versatile and robust centrifugation-based filtration protocol for the isolation and quantification of α-synuclein monomers, oligomers and fibrils: Towards improving experimental reproducibility in α-synuclein research. J. Neurochem. 2020, 153 (1), 103−119.

20 Dikiy, I.; Fauvet, B.; Jovicic, A.; Mahul-Mellier, A. L.; Desobry, C.; El-Turk, F.; Gitler, A. D.; Lashuel, H. A.; Eliezer, D. Semisynthetic and in Vitro Phosphorylation of Alpha-Synuclein at Y39 Promotes Functional Partly Helical Membrane-Bound States Resembling Those Induced by PD Mutations. ACS Chem. Biol. 2016, 11 (9), 2428−2437.

21 Donzelli, S., OSullivan, S. A., Mahul-Mellier, A. L., Ulusoy, A., Fusco, G., Kumar, S. T., & Lashuel, H. A. (2023). Post-fibrillization nitration of alpha-synuclein abolishes its seeding activity and pathology formation in primary neurons and in vivo. bioRxiv, 2023, 10.1101/2023.03.24.534149

22 (a) Kim, Y.; Vaidya, B.; McInnes, J.; Zoghbi, H. Y. Alpha-Synuclein Phosphomimetic Y39E and S129D Knock-In Mice Show Cytosolic Alpha-Synuclein Localization without Developing Neurodegeneration or Motor Deficits. eneuro, 2025, 12(4). (b) Zhang, S.; Zhu, R.; Pan, B.; et al. Post-translational modifications of soluble α-synuclein regulate the amplification of pathological α-synuclein. Nat Neurosci 2023, *26*, 213–225.

23 (a) Mahul-Mellier, A. L.; Bruno Fauvet, Amanda Gysbers, Igor Dikiy, Abid Oueslati, Sandrine Georgeon, Allan J. Lamontanara, et al. "c-Abl phosphorylates α-synuclein and regulates its degradation: implication for α-synuclein clearance and contribution to the pathogenesis of Parkinson’s disease." Human molecular genetics. 2014, 23 (11), 2858–2879. (b) Brahmachari, S,; Ge, P.; Lee, S. H.; Kim, D.; Karuppagounder, S. S.; Kumar, M.; Mao, X.; et al. Activation of tyrosine kinase c-Abl contributes to α-synuclein– induced neurodegeneration. J Clin Invest. 2016, 126 (8), 2970–2988.

24 (a) Shults, C. W., Lewy bodies. Proc. Natl. Acad. Sci. U.S.A. 2006, 103, 1661–1668. (b) Shahmoradian, S. H. et al., Lewy pathology in Parkinson’s disease consists of crowded organelles and lipid membranes. Nat. Neurosci. 2019, 22, 1099–1109.

25 (a) Chen, L., et al. Tyrosine and serine phosphorylation of α-synuclein have opposing effects on neurotoxicity and soluble oligomer formation. J. Clin. Investig. 2009, 10.1172/JCI39088. (b) Altay, M. F.; Kumar, S. T.; Burtscher, J.; et al. Development and validation of an expanded antibody toolset that captures alpha-synuclein pathological diversity in Lewy body diseases. npj Parkinsons Dis. 2023, 9, 161.

26 Hejjaoui, M.; Butterfield, S.; Fauvet, B.; Vercruysse, F.; Cui, J.; Dikiy, I.; Prudent, M.; Olschewski, D.; Zhang, Y.; Eliezer, D.;, et al. Elucidating the Role of C-Terminal Post-Translational Modifications Using Protein Semisynthesis Strategies: α-Synuclein Phosphorylation at Tyrosine 125. J. Am. Chem. Soc. 2012, 134 (11), 5196−5210.

27 Oueslati, A.; Fournier, M.; Lashuel, H. A. "Role of post-translational modifications in modulating the structure, function and toxicity of α-synuclein: implications for Parkinson’s disease pathogenesis and therapies." Progress in Brain Research, 2010, 183, 115–145.

28 (a) Nakamura, T.; Yamashita, H.; Takahashi, T.; Nakamura, S. Activated Fyn phosphorylates α-synuclein at tyrosine residue 125. Biochemical and biophysical research communications, 2001, 280 (4), 1085–1092. (b) Ellis, C. E.; Schwartzberg, P. L.; Grider, T. L.; Fink, D. W.; Nussbaum, R. L. alpha-synuclein is phosphorylated by members of the Src family of protein-tyrosine kinases. J Biol Chem. 2001, 276 (6), 3879–84. (c) Negro, A., Brunati, A. M., Donella-Deana, A., Massimino, M. L.; Pinna, L. A. (2002). Multiple phosphorylation of α-synuclein by protein tyrosine kinase Syk prevents eosin-induced aggregation. The FASEB Journal, 2002, 16 (2), 1–22.

29 Gatzemeier, L. M.; Meyer, F.; Outeiro, T. F. Synthesis and Semi-Synthesis of Alpha-Synuclein: Insight into the Chemical Complexity of Synucleinopathies. ChemBioChem. 2024, 25 (20), e202400253.

30 Zhao, K.; Lim, Y.; Liu, Z.; Long, H.; Sun, Y.; Hu, J.; Zhao, C.; Tao, Y.; Zhang, X.; Li, D.; Li, Y.; Liu, C. Parkinson’s disease-related phosphorylation at Tyr39 rearranges α-synuclein amyloid fibril structure revealed by cryo-EM. Proc. Natl. Acad. Sci. U.S.A. 2020, 117 (33), 20305–20315.

