## Supplementary material for "A Site-Specific Organometallic Approach for Installing Tyrosine Phosphorylation Mimics to Decipher the Role of Phosphorylation in Alpha-Synuclein (aSyn) Aggregation and Seeding": The supporting information includes synthesis and methods, as well as biochemical, biophysical and biological characterization of all compounds.

### Table of contents


1. Experimental …………………………………………………………………..4-5
2. Synthesis of the palladium(II) oxidative addition complex (Pd-OAC-NP) …………5-6
3. Late-stage transfer of Tyr-phosphorylation mimic using Pd-OAC-NP and stability of the final construct ..…………………………………………………………...7-12
4. Preparation and characterization of recombinant aSyn species …………………..13-14
5. General biochemical, biophysical, and biological experimental methods …….....14-19
6. List of antibodies for dot-blot (table S1) ……………………………………………16
7. List of antibodies for ICC (table S2) ………………………………………………..18-19
8. UPLC-MS traces of aSyn Cys-mutants ……………………………………………..19-20
9. Site-specific aSyn phosphorylation via Pd-OAC-NP ……………………...……20-28
10. SDS-PAGE of all the aSyn phosphorylated analogues ………………………………29
11. Dot-blot and phosphatase assays …………………………………………………...30
12. Aggregation ThT assay [WT vs. mono-P(nat) analogues] …………………………...31
13. Sedimentation Assay for aSyn WT, mono-P(nat), di-P(nat) and tri-P(nat) post-aggregation …………………………………………………………………………..32
14. Characterization of the fibrils of aSyn WT, mono-P(nat), di-P(nat) and tri-P(nat) post-aggregation …………………………………………………………………………..33
15. Generation of different aSyn PFF species (purity and characterization) ……………………………………………………………………………..………34
16. Generation of WT aSyn (h) stably expressing U2OS cell line ………………………35
17. pS129 pathology in PFF treated WT-aSyn (h) stably-expressing U2OS cells ……………………………………………………………………………………….36-37
18. NMR spectra of synthesized compounds…………………………………………….38-40
19. References...……………………………………………………………………………41

#### 1. Experimental

##### 1.1 Materials

Fmoc-L-Phe-OH, Fmoc-L-Asn(Trt)-OH, Fmoc-L-Gln(Trt)-OH, Fmoc-L-Arg(Pbf)-OH, Fmoc-L-Tyr(tBu)-OH, Fmoc-L-Glu(OtBu)-OH, Fmoc-L-Ala-OH, Fmoc-L-Leu-OH, Fmoc-L-His(Trt)-OH, Fmoc-L-Asp(OtBu)-OH, Fmoc-L-Pro-OH, Fmoc-L-Cys(Trt)-OH, Fmoc-L-Lys(Boc)-OH, Fmoc-L-Lys(Alloc)-OH, Fmoc-L-Ile-OH, Fmoc-L-Ser(tBu)-OH, Fmoc-Gly-OH, Fmoc-L-Nle-OH, Fmoc-L-Thr(tBu)-OH, Boc-L-Ala-OH, Boc-L-asparagine, 1,3-Diisopropylcarbodiimide (DIC), 4-Bromobenzylphosphonic acid were purchased from Sigma-Aldrich. Fmoc-L-His(Boc)-OH was purchased from CEM. 1 [Bis(dimethylamino)methylene]-1H-1,2,3-triazolo[4,5-b] pyridinium 3-oxide hexafluorophosphate (HATU), (2-(1H-benzotriazol-1-yl)-1,1,3,3-tetramethyluronium hexafluorophosphate (HBTU), and HOBt HYDRATE were purchased from Luxembourg Bio Technologies Ltd. Rink Amide ProTide resin was obtained from CEM. 2 Dicyclohexylphosphino-2′,6′-diisopropoxybiphenyl (RuPhos) was purchased from Angene Chemical Private Limited. 4-Bromo-2-nitrophenol (98%) was purchased from Thermo Scientific. Diethyl ether ($Et_2O$, 99.8% stabilized, ACS grade) was obtained from MACRON. Dichloromethane ReagentPlus®, diisopropylethylamine (DIEA, ReagentPlus®), triisopropylsilane (TIS, 98%), formic acid (FA, 98-100% for LC/MS), n-Pentane (AR) were purchased from Bio-Lab Ltd. Water for all reactions carried out on proteins and for reverse-phase purification was obtained via filtration of deionized water through a MilliporeSigmaTM Milli-QTM Ultrapure Water System. All chemicals obtained from the supplier were used as received without further purification. ($CH_2Cl_2$, ≥99.5% stabilized with 50 ppm Amylene) was obtained from CHEM-LAB. N, N- dimethylformamide (DMF, Peptide Synthesis-grade) and acetonitrile (ACN, LC/MS Grade) were purchased from J.T.Baker. Trifluoroacetic acid (TFA, ≥99% ≥99% ReagentPlus®), piperidine (≥99%

##### 1.2 General Fmoc-SPPS procedure

Fmoc-SPPS was carried out in the presence of 4 eq. of AA, 4 eq. of HCTU, and 8 eq. of DIEA to the initial loading of the resin for 50 min. The dipeptides were coupled manually using 2.5 eq. of AA, 2.5 eq. of HATU, and 5 eq. of DIEA to the initial loading of the resin for 1 h 30 min. To cleave the peptides from the solid support, the resin was washed with DMF, MeOH, DCM (X5), and dried under high vacuum. A cocktail of TFA: triisopropyl silane (TIS): water (95:2.5:2.5) was added to the resin, and the reaction mixture was shaken for 2 h at RT. The resin was filtered, and the combined filtrate was added dropwise to a 10-fold volume of cold ether, which was then centrifuged. The precipitate was dissolved in acetonitrile-water for freeze-drying in the lyophilizer, yielding the crude peptide.

#### 1.3 Liquid chromatography mass spectrometry (LC-MS) analysis

Analytical RP-HPLC chromatograms were acquired using Thermo Scientific Vanquish HPLC. Method A: XBridge® BEH 3.5 µm-C4 300 Å Column (150 x 4.6 mm); LC conditions: 5% B from 0–1.0 min, then a linear gradient from 5% to 60% B from 1.0–28.0 min, 1.0 mL/min flow rate. Mobile phases used are solvent A (0.05% TFA in water) and solvent B (0.05% TFA in acetonitrile).

Method B: BioZen™ 2.6 µm-C4 Widepore LC column (150 x 2.1 mm); LC conditions: 5% B from 0–1.0 min, then a linear gradient from 5% to 50% B from 1.0–11.0 min, 0.3 mL/min flow rate. Mobile phases used are solvent A (0.1 % FA in water) and solvent B (0.1 % FA in acetonitrile).

Mass spectra were acquired using a Thermo Scientific ISQ EM Mass spectrometer.

#### 1.4 Preparative RP-HPLC purification

Preparative RP-HPLC was performed using Thermo Scientific DIONEX UltiMate 3000 Variable Wavelength Detector, equipped with a column of choice. Mobile phases used for LC analysis were solvent A (0.05% TFA in water) and solvent B (0.05% TFA in acetonitrile). The following LC methods were used:

<u>Method A:</u> XBridge® Protein BEH C4 OBD™ Prep Column, 300 Å, 5 µm, 250 mm x 10 mm, 1/pkg, LC conditions: 5% B from 0–5.0 min, then linear gradient from 5% to 60% B from 5-65 min, 4 mL/min flow rate at 30 °C.

<u>Method B:</u> Jupiter® 5 µm C4 300 Å LC column (250 x 10 mm), LC conditions: 5% B from 0–5.0 min, then linear gradient from 5% to 60% B from 5-65 min, 4 mL/min flow rate at 30 °C.

#### 2. Synthesis of the palladium(II) oxidative addition complex (Pd-OAC-NP)

#### 2.1 Synthesis of 4-iodophenyl dihydrogen phosphate

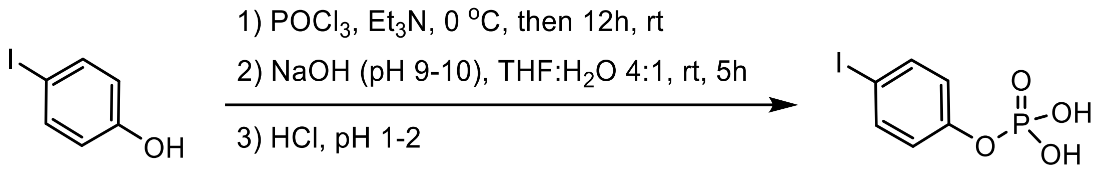


4-iodophenol (660 mg, 3 mmol, 1 equiv.) was dissolved in distilled ether (2 mL) in a flask equipped with a stir bar. The reaction solution was cooled down to 0 °C. $POCl_3$ (279 µL, 3 mmol, 1 equiv.) was gradually added. Then $Et_3N$ (419 µL, 3 mmol, 1 equiv.) was slowly added and kept for 1h.[1] The reaction was allowed to stir at room temperature for 12 h. Subsequently, the reaction mixture was filtered, and the filtrate was evaporated under vacuum to afford 4-iodophenyl dihydrogen phosphate as a yellow liquid. Without further purification, 4-iodophenyl dihydrogen phosphate was dissolved in THF: $H_2O$ (4:1) solution, and the pH was adjusted to 9-10 using 5 *N* NaOH and stirred at room temperature for 5h to form sodium 4-iodophenyl phosphate. The solution was acidified to pH 1-2 using 5 *N* HCl. 1h later, the solution was diluted with $H_2O$ and purified by flash chromatography using the Biotage® Selekt system, affording the product (335 mg, 37.2%) as a white powder.

The purification was performed using reverse phase (RP) chromatography via Biotage® Selekt using a 25 g Biotage Sfär Bio C18 D20 µm column. The mobile phases used in the flash purification were solvent A (0.1% TFA in water) and solvent B (0.1% TFA in acetonitrile). The gradient elution conditions were as follows: 5% B for 2 column volumes (CV), a linear ramp from 5% to 95% B over 10 column volumes (CV), 90% B for 2 column volumes (CV), at a flow rate of 40 mL/min.

$^{1}$H NMR (400 MHz, MeOD) δ 7.65 (d, $J$ = 8.4 Hz, 2H), 7.01 (d, $J$ = 8.0 Hz, 2H). $^{31}$P NMR (162 MHz, MeOD) δ -3.70 (s). HRMS (ESI) m/z: $[M-H]^-$ calculated for $[C_6H_5IO_4P]^-$ 298.8970, found 298.8972.

#### 2.2 Synthesis of Pd-OAC-NP:

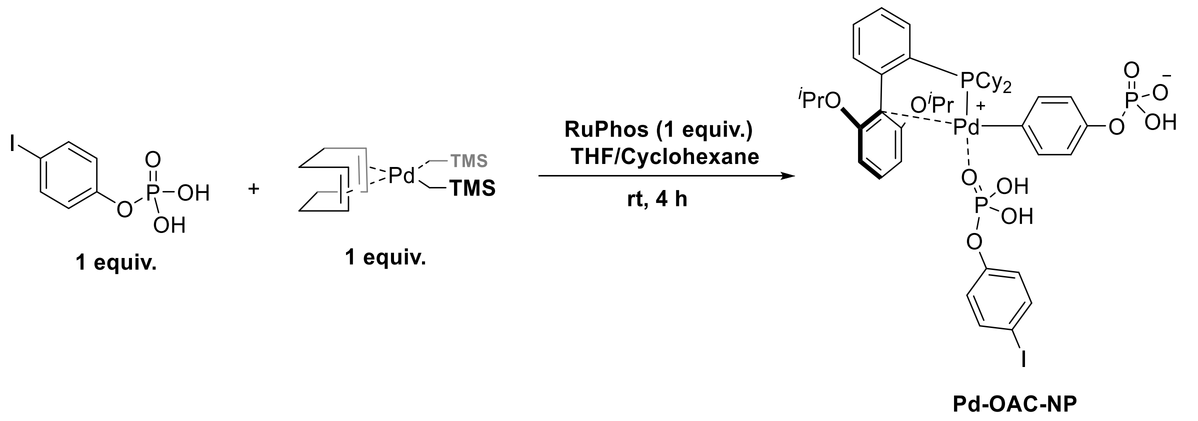


In a 50 mL Schlenk flask equipped with a magnetic stir bar, RuPhos (23.3 mg, 0.05 mmol, 1.0 equiv.) and 4-iodophenyl dihydrogen phosphate (15 mg, 0.05 mmol, 1.0 equiv.) were dissolved in 1 mL dry THF.[2] Solid (COD)Pd($CH_2SiMe_3$)$_2$ (19.5 mg, 0.05 mmol, 1.0 equiv.) was dissolved in Cyclohexane (1 mL) added rapidly in one portion. The resulting solution was stirred for 4 h at room temperature. After that, cold pentane was added, and the resulting mixture was transferred to a vial (20 mL) and placed into a –20 °C freezer for 3 h. The vial was then taken outside of the freezer and the resulting precipitate was collected, washed with pentane (3 × 3 mL), and dried under vacuum to afford **Pd-OAC-NP** as yellow powder. (26.1 mg, yield 50 %)

$^{1}$H NMR (400 MHz, $CD_2Cl_2$) δ 7.55 (d, *J*= 0.5 Hz, 3H), 7.42 (dd, *J* = 15.3, 7.1 Hz, 2H), 7.13 (s, 2H), 6.94 (s, 4H), 6.80 (s, 2H), 6.64 (s, 1H), 6.45 (s, 1H), 4.54 (s, 2H), 2.16 (s, 2H), 1.74 (d, *J* = 38.1 Hz, 12H), 1.43 (s, 4H), 1.20 (d, *J* = 40.3 Hz, 9H), 1.03 (s, 6H), 0.83 (s, 1H).$^{31}$P NMR (162 MHz, DMF-$d_7$) δ 46.0 (s), -2.1 (s). $^{13}$C NMR (101 MHz, $CD_2Cl_2$) δ 160.08 (s), 157.97 (s), 152.28 (s), 149.57 (s), 144.84 (s), 137.98 (s), 136.91 (d, *J* = 45.3 Hz), 131.77 (d, *J* = 48.8 Hz), 130.77 (d, *J* = 33.2 Hz), 126.19 (s), 122.51 (s), 120.39 (s), 107.10 (d, *J* = 23.1 Hz), 106.46 (s), 105.52 (s), 86.30 (s), 70.68 (s), 35.11 (d, *J* = 29.2 Hz), 34.20 (d, *J* = 30.2 Hz), 29.03 (d, *J* = 37.1 Hz), 28.03 (s), 27.07 – 26.49 (m), 25.87 (s), 21.45 (s). HRMS (ESI): m/z: calcd. for [$C_{42}H_{54}O_{10}P_3IPd^{[104]}$-H]$^-$: 1041.0936, found : 1041.0937.

#### 3. Late-stage transfer of Tyr-phosphorylation mimic using Pd-OAC-NP

##### 3.1 Modification of model peptide P1 with Pd-OAC-NP

P1 was synthesized as described previously,[4] and the reaction was carried out according to the following scheme:

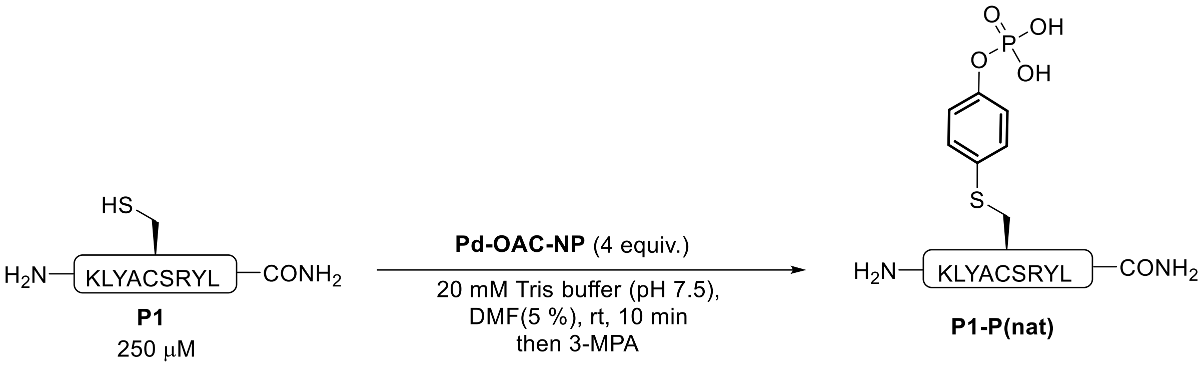


To a 1.5 mL Eppendorf was added P1 (937 μL, 26 μM, 1.0 equiv.) in a 20 mM Tris (pH 7.5) solution and Pd-OAC-NP (49 μL, 20 mM, 4.0 equiv.) as a solution in DMF (v = 5 %).[3,4] The final reaction concentrations of the major reaction components were as follows: P1 (0.25 mM) and Pd-OAC-NP (1.0 mM). The reaction mixture was vigorously stirred for a few seconds and then kept at room temperature for 10 min. Next, the Eppendorf was opened, and the reaction was treated with 3-MPA solution (0.2 μL, 10 equiv. compared to P1) and was kept at room temperature for 10 min. 10 μL of the reaction mixture was injected into the LC-MS using method A.

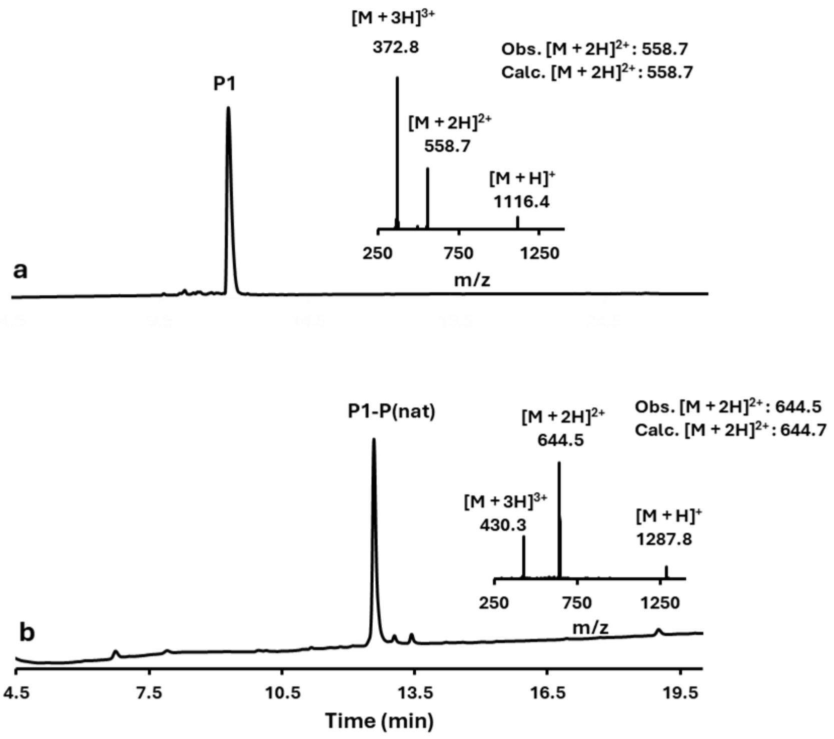


**Figure S1. LCMS analysis of the phosphorylation of model peptide P1 using Pd-OAC-NP.** LC chromatogram of the UV absorbance at 214 nm and mass-to-charge (m/z) spectra. (a) peptide P1 at t = 0. (b) crude phosphorylation reaction of P1 with Pd-OAC-NP at t = 10 min.

##### 3.2 Modification of model protein MaxA61C via Pd-OAC-NP

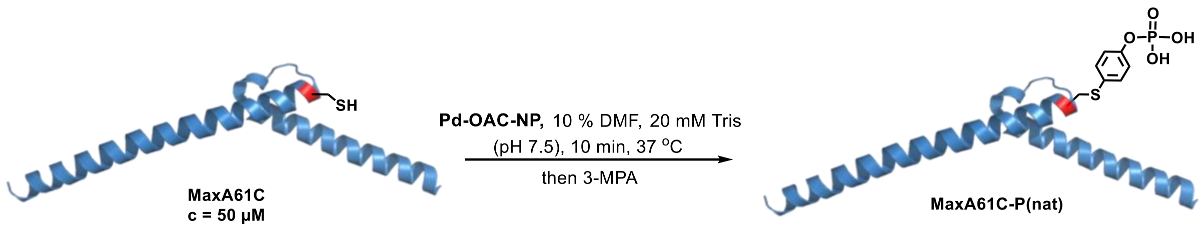


To a 1.5 mL Eppendorf was added fresh prepared **MaxA61C** Tris solution (93 µL, 55 µM, 1.0 equiv.) and the **Max** solution was prewarmed at 37 °C for 1 min. After that, **Pd-OAC-NP** (5.2 µL, 7.5 mM, 7.5 equiv.) was diluted with 5.2 µL pure DMF and then was added to Max solution in one portion (v = 10 %). The final reaction concentrations of the major reaction components were as follows: **MaxA61C** (50 µM) and **Pd-OAC-NP** (375 µM). The reaction mixture was mixed by pipetting up and down 10× followed by a 10 min incubation at 37 °C. Next, the Eppendorf was opened and the reaction was treated with 3-MPA solution (1 µL from 4.5 µL pure 3-MPA in 995.5 µL 20 mM Tris (pH 7.5), 10 equiv. compared to Max) and was kept at room temperature for 10 min. 10 µL of the reaction mixture was injected into the LC-MS using method B.

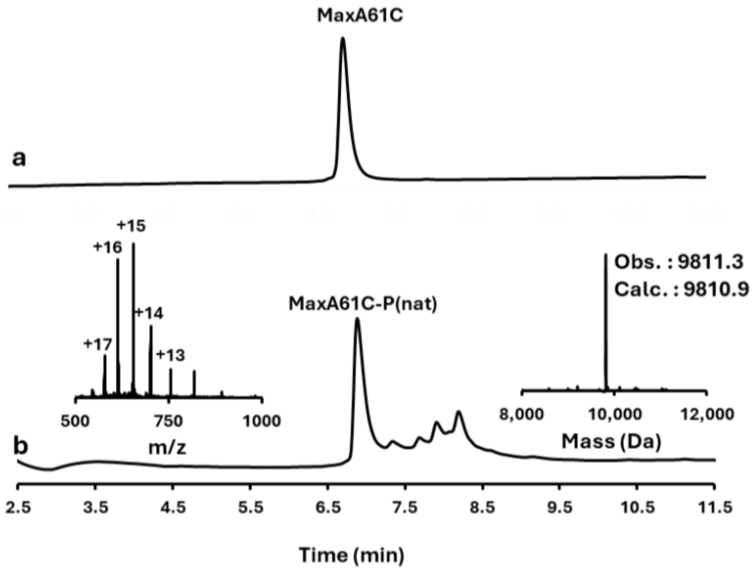


**Figure S2. LCMS analysis of MaxA61C phosphorylation via Pd-OAC-NP.** LC chromatogram of the UV absorbance at 214 nm, mass-to-charge (m/z) spectra, and deconvoluted mass spectra. (a) **MaxA61C** at t = 0. (b) crude phosphorylation reaction of **MaxA61C** with **Pd-OAC-NP** at t = 10 min.

#### 4. Isolation of modified MaxA61C-P(nat) analog and stability analysis

##### 4.1 Isolation of native phosphorylated MaxA61C-P(nat)

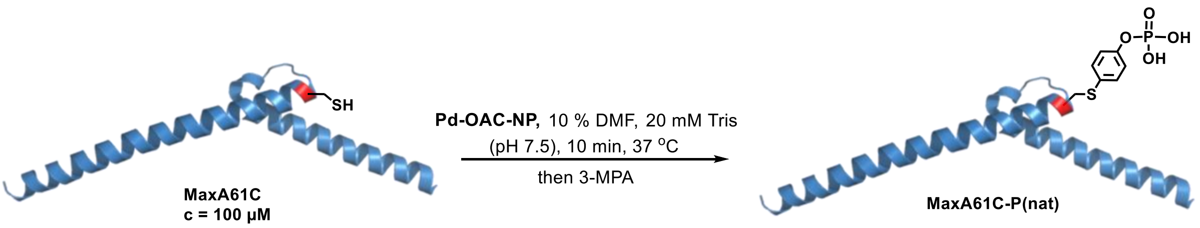


To a 15 mL falcon tube was added **MaxA61C** (3.9 mg, 3.6 mL, 111 µM, 1.0 equiv.) in 20 mM Tris (pH 7.5) solution and prewarmed at 37 °C for 1 min, followed by quick addition of **Pd-OAC-NP** (3.2 mg, 404 µL, 7.5 mM, 7.5 equiv.) as a solution in DMF (v = 10 %). The final reaction concentrations of the major reaction components were as follows: **MaxA61C** (100 µM) and **Pd-OAC-NP** (750 µM). The reaction mixture was vigorously stirred for a few seconds and followed by a 10 min incubation at 37 °C. Next, the falcon tube was opened, and the reaction was treated with 3-MPA (0.35 µL, 10 equiv. compared to Max) and was kept at room temperature for 10 min. 10 µL of the reaction mixture was injected into the LC-MS using method B. The phosphorylated product was purified by RP-HPLC by using Method A, affording 2.3 mg as a white powder. (isolated yield 56 %). $^{31}$P NMR (162 MHz, $D_2O$) δ -2.90 (s).

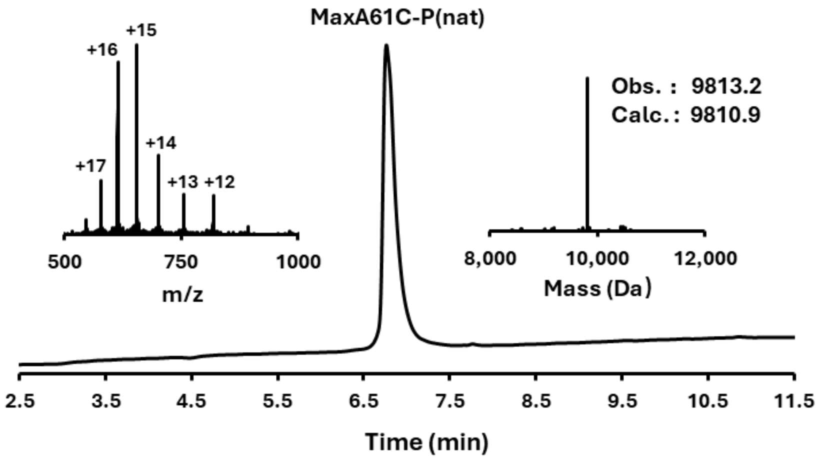


**Figure S3. LCMS analysis of the isolated MaxA61C-P(nat).** LC chromatogram of the UV absorbance at 214 nm, mass-to-charge (m/z) spectrum, and deconvoluted mass spectrum.

#### 4.2 Stability of isolated native Phosphorylated MaxA61C-P(nat)

Isolated MaxA61C-P(nat) (0.5 mg) was dissolved in PBS buffer (x1) (255 µL, C = 200 µM, pH 7.0), and the solution was analyzed by LC-MS using method B. The solution was incubated at 37 °C. The stability reaction was monitored by LC-MS using the same method after 1 day, 3 days, and 5 days.

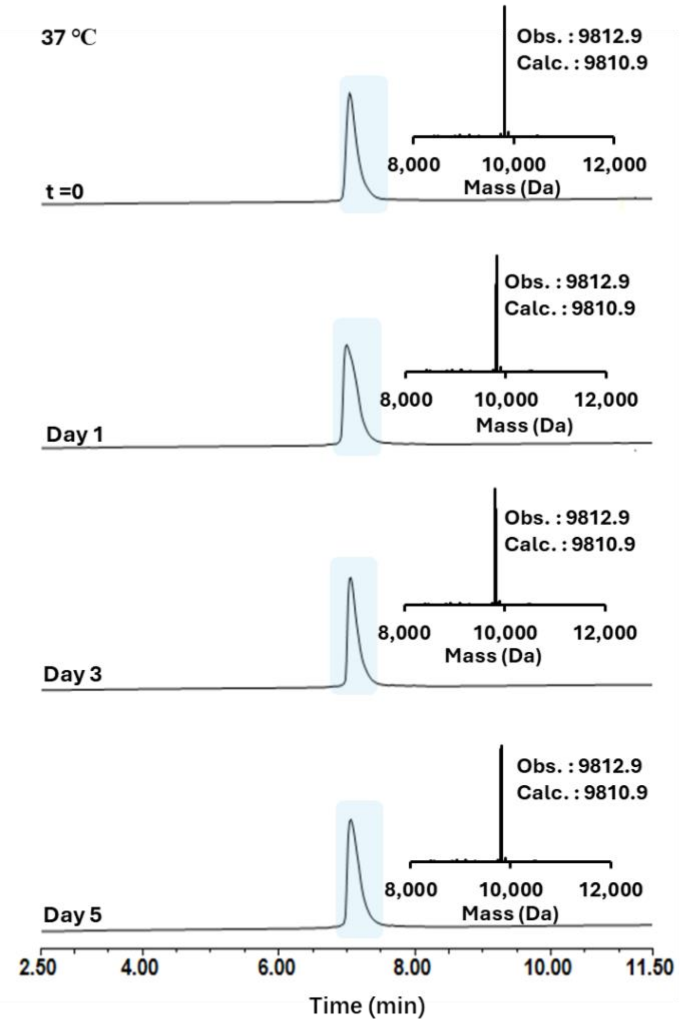


**Figure S4. LCMS analysis of the stability of isolated MaxA61C-P(nat).** LC chromatogram of the UV absorbance at 214 nm, mass-to-charge (m/z) spectrum, and deconvoluted mass spectrum.

#### 5. Preparation and characterization of recombinant aSyn species:

##### 5.1 Amino acid Sequence of human WT aSyn:

MDVFMKGLSKAKEGVVAAAEKTKQGVAEAAGKTKEGVL**Y**$^{39}$VGSKTKEGVVHGVA TVAEKTKEQVTNVGGAVVTGVTAVAQKTVEGAGSIAAATGFVKKDQLGKNEEGAPQ EGILEDMPVDPDNEA**Y**$^{125}$EMPSEEG**Y**$^{133}$QD**Y**$^{136}$EPEA

*Highlighted phosphorylation sites (Tyr 39, 125, 133,136)*

BL21 (DE3) cells transformed with the pT7-7 plasmid encoding the WT aSyn or Y39C or Y125C or Y133C or Y136C or Y125C/Y133C or Y125C/Y133C/Y136C aSyn mutants were grown in LB medium at 37°C and induced with 1 mM 1-thio-β-d-galactopyranoside (AppliChem) at an O.D. between 0.4–0.6 and continued to grow for 4 h at 37°C. Following the overnight incubation at 18 °C, induced bacterial cultures were pelleted and sonicated for cell lysis. After centrifugation at 18,000 g for 20 min, the supernatant was boiled for 5 min and centrifuged again for 20 min. The supernatant was purified by anion exchange chromatography (HiPrep 16/10 Q FF, GE Healthcare Life Sciences), followed by reverse-phase HPLC (Jupiter 300 C4, 20 mm I.D. × 250 mm, 10 μm average bead diameter, Phenomenex) and lyophilized. The WT, mono-Cys mutants of aSyn were used from the previous expression from reference 5.

##### 5.2 Mass spectrometry analysis:

Mass spectrometry (MS) analysis of proteins was performed by liquid chromatography-mass spectrometry (LC-MS) on the LTQ system (Thermo Scientific, San Jose, CA). Before analysis, proteins were desalted online by reversed-phase chromatography on a Poroshell 300SB C3 column (1.0 x 75 mm, 5 µm, Agilent Technologies, Santa Clara, CA, on the LTQ system). 10 µL protein samples were injected into the column at a flow rate of 300 uL/min and were eluted from 5% to 95% of solvent B against solvent A, linear gradient. The solvent composition was Solvent A: 0.1% formic acid in ultra-pure water; Solvent B: 0.1% formic acid in acetonitrile. MagTran software (Amgen Inc., Thousand Oaks, CA) was used for charge state deconvolution and MS analysis.

##### 5.3 SDS-PAGE analysis:

Samples for SDS-PAGE were mixed with 2 × Laemmli buffer (4% SDS, 20% glycerol, 0.004% bromphenol blue, 0.125 M Tris-Cl, 10% 2-mercaptoethanol, pH 6.8) and loaded onto 16% polyacrylamide tricine gels. The gel was run at 120 V for 2 hr in running buffer (25 mM Tris,

192 mM Glycine, 0.1% SDS, pH 8.3), followed by staining with a solution of 25% (v/v) isopropanol, 10% acetic acid (v/v) and 0.05% (w/v) or commertially available Coomassie brilliant blue R (Applichem) and destaining with boiling distilled water.

#### 6. Transmission electron microscopy (TEM):

Before the application of the sample, Formvar and carbon-coated 200 mesh-containing copper EM grids (Electron Microscopy Sciences) were glow-discharged for 30 s at 20 mA using a PELCOeasiGlow™ Glow Discharge Cleaning System (TED PELLA, Inc). Subsequently, 5 µL of the sample was placed onto the EM grid for a minute. Then, the sample was carefully blotted using filter paper and air-dried for 30 s. Then, the grids were washed three times with ultrapure water and stained with 1% (w/v) uranyl formate solution. Grids were examined using a Bio Talos electron microscope. The microscope was equipped with a LaB6 gun operated at an acceleration voltage of 80 kV, and images were captured using a 4K × 4K charge-coupled device camera (FEI Eagle). The length (ln) or diameter (d) of respective aSyn oligomeric fractions was measured using ImageJ software (NIH).

#### 7. ThT Aggregation Kinetics Assay:

Lyophilized WT, Cys mutants, phosphorylated, were dissolved to about 30 µM in PBS buffer (pH 7.4). Protein solutions were then filtered through 100 kDa MWCO Microcon filters to remove any insoluble material or oligomers at the beginning of the experiment. Protein concentrations were precisely measured by UV spectroscopy (using extinction coefficients of ε280nm = 5960 $M^{-1}cm^{-1}$ for wild-type aSyn and ε280nm = 4470 $M^{-1}cm^{-1}$ for mutant and modified aSyn). Proteins were then diluted to a final concentration of 15 or 20 µM (final volume: 400 µL). A fresh ThT solution (0.01 mM ThT solution in 0.5 M glycine, pH 8.5) was added to each (a total of 13) protein sample, and 100 µL solution was added to a clear-bottom 96-well plate (Corning). The plate was then sealed with a clear film. Aggregations were conducted in the FLUstar Omega microplate reader (BMG Labtech Germany) at an excitation wavelength of 450 nm and an emission wavelength of 485 nm. The reading was taken from time 0 to every 8 min for the next 3 days with ideal shaking at 37°C. Three independent experiments were performed in triplicate for each protein.

#### 8. Sedimentation assay:

Our well-established centrifugation-based assay can separate sedimentable fibrils from the monomers and oligomers. First, 10 µL of the sample after aggregation was taken out and diluted with 2X LB. This is represented as "total". After, a total sample volume of 50 µL was subjected to ultracentrifugation at 100,000 g for 30 min at 4°C. Following centrifugation, careful pipetting of the supernatant from the pellet is carried out to isolate the soluble species, which represents the monomers & oligomers (supernatant). The isolated pellet was resuspended in 50 µL PBS and represented as Fibrils (pellet). These three fractions from all 13 protein samples were run in 16% tricine gels and stained with Instant Blue Coomassie solution to visualize the distribution.

#### 9. Dot-Blot analysis:

Nitrocellulose membranes (Amersham, 10,600,001) were spotted with 5 µL of 20 µM protein samples of aSyn species. The membranes were blocked for 1 h with Odyssey blocking buffer (LiCoR, 927-40000) and then incubated overnight with different primary antibodies (Table S1) diluted in PBS at 1:500 dilution at 4 ºC. The membranes were washed three times, with 0.1% PBS-Tween (3 × 10 minutes) and incubated with IR dye-conjugated secondary antibodies (1:10000) (Table S1) for 1h at RT. The membranes were washed three times, with 0.1% PBS-Tween (3 × 10 minutes). The visualization was performed by fluorescence using Odyssey CLx from LiCor.

#### 10. Lambda protein phosphatase (LPP) assay:

All the reagents were supplied along with the LLP kit purchased from New England Biolabs, and aSynY39C-P(nat) and aSynY125C-P(nat) were reacted with Lambda Protein Phosphatase (LLP) as per standard protocol. Briefly, the protein samples were dissolved in PBS (20 µM) to a total volume of 40 µL, and to this, 5 µL of 10X NEBuffer for Protein Metallo Phosphatases (PMP) and 5 µL of 10 mM $MnCl_2$ were added, making a total reaction volume of 50 µL. Then 1 µl of Lambda Protein Phosphatase was added and incubated at 37 °C for 2 h. The phosphatase activity was tested using the site-specific PTM-dependent aSyn antibodies via the dot blot method as described in the previous section.

**Table S1: Primary and secondary antibodies used in Dot-Blot and phosphatase experiments.**

| Primary Antibody | Name | Species | Epitope | Dilution |
|---|---|---|---|---|
| | Anti-aSyn (SYN-1) Code : B59 | Mouse Monoclonal | 91-99 | 1:1000 |
| | Anti aSyn pY39 Code : B65 | Rabbit Polyclonal | pY39 | 1:500 |
| | Anti aSyn pY125 Code : B69 | Rabbit Polyclonal | pY125 | 1:1000 |
| | Anti aSyn pY133 Code : B291 | Rabbit Polyclonal | pY133 | 1:500 |
| | Anti aSyn pY136 Code : B292 | Rabbit Polyclonal | 134-138; pY136 | 1:500 |
| Secondary Antibody | Goat anti-Rabbit Alexa Flour 800nm | - | - | 1:10000 |
| | Goat anti Mouse Alexa Flour 680 nm | - | - | 1:10000 |

#### 11. Preparation of WT, Y39C and Y39C-P(nat) aSyn monomers:

The lyophilized aSyn of the required quantity was resuspended in PBS buffer and followed by the filtration protocol as described before.[5] Briefly, the lyophilized aSyn following resuspension was adjusted to pH ~7.2 to 7.4 and filtered using 100-kDa spin filters. The filtrate containing monomers was estimated for the concentration by measuring the absorbance at 280 nm using a NanoDrop 2000 spectrophotometer (Thermo Fisher Scientific) and using an extinction coefficient of 5960 $M^{-1}$ $cm^{-1}$ predicted using the sequence (ProtParam, ExPASy). Monomeric aSyn (WT, Y39C, and Y39C-P(nat)) was prepared as described above, followed by any kind of further experiments

#### 12. Preparation of WT, Y39C and Y39C-P(nat) aSyn PFF seeds for cell studies:

WT, Y39C, and Y39C-P(nat) aSyn fibrils were prepared by dissolving 4 mg of lyophilized recombinant aSyn in 600 µl of 1× PBS, and the pH was adjusted to ~7.2 to 7.4. The solution was filtered through 0.2-µm filters (Merck, SLGP033RS), and the filtrate was transferred to black screw cap tubes. This tube was incubated in a shaking incubator at 37°C and 1000 rpm for 5 days. After 5 days, the formation of fibrils was assessed by TEM and Coomassie staining as described previously.[5] For the preparation of sonicated seeds, WT, Y39C, and Y39C-P(nat) aSyn fibrils were subjected to sonication for 20 s, at 20% amplitude, 1-s pulse on, and 1-s pulse off (Sonic Vibra-Cell, Blanc-Labo, Switzerland) only in mentioned experiments. The amount of monomers and oligomers released from sonicated fibrils was quantified by filtration following the published protocol.[5] The structure of the fibrils was monitored by TEM and the number of released monomers and oligomers after sonication was determined using Coomassie staining analysis. The length of the sonicated PFFs was calculated using ImageJ software.

#### 13. *In vitro* cross-seeding monitoring via ThT aggregation assay:

For the cross-seeding aggregation experiments, 1% WT or Y39C or Y39C-P(nat) PFFs (w/v, relative to the monomer concentration) were added to the 400-µl master mix solution containing the final concentration (20 µM) of WT aSyn monomer in 1× PBS, to which a final ThT concentration of 8 µM was added. For all the ThT-based aggregation assays, 100 µl of replicates from each master mix solution was added to the three individual wells in a black 96-well optimal bottom plate (Costar), with each well pre-added with six SiLibeads ceramic beads (Sigmund Lindner) with a diameter of 1.0 to 1.2 mm. The plate was sealed with Corning microplate tape and transferred to a FLUOstar OPTIMA plate reader (BMG Labtech). The parameters used were as follows: continuous orbital shaking at 600 rpm at 37 °C; ThT fluorescence was monitored every 8 s using excitation at 450 ± 10 nm and emission at 480 ± 10 nm. At the end point of the assays, the samples were analyzed by EM.

#### 14. Mammalian Cell culture:

U2OS cells were cultured at 95% air and 5% $CO_2$ in Dulbecco's modified Eagle's medium (DMEM; low glucose) supplemented with 10% fetal bovine serum (Gibco) and penicillin-streptomycin (Thermo Fisher Scientific).

#### 15. Preparation and treatment of WT hippocampal primary neurons:

WT primary hippocampal neurons were prepared from WT C57BL/6JRj (Janvier) pups at postnatal day 0 (P0), as previously reported.[6] The dissociated neurons were plated in 6-well plates, onto coverslips (VWR) or in clear black-bottom 96-well plates (Falcon) freshly coated with poly-l-lysine 0.1% wt/vol in water (Brunschwig). Neurons were plated at a density of 300,000 cells per ml for biochemistry analyses, 250,000 cells per ml for the immunocytochemistry (ICC) analyses, and 200,000 cells per ml for the high-content imaging analysis (HCA). In WT hippocampal primary neurons, the PFF-treated neurons were all fixed at DIV 14 post-treatment. PBS was used as a negative control. All procedures were approved by the Swiss Federal Veterinary Office (authorization number VD3496).

#### 16. Immunocytochemistry (ICC):

Immortal cells (U2OS stably expressing WT aSyn (h)) or WT primary hippocampal neurons were washed twice with PBS, fixed in 4% paraformaldehyde (PFA) for 20 min at RT, and immunostained as previously described (in ref. 6). The antibodies used are indicated in the corresponding legend of each figure. The source and dilution of each antibody are listed in Supplementary Table S2. The cells plated in black, clear-bottom, 96-well plates were imaged using the IN Cell Analyzer 2200 (with a 10× objective). For each independent experiment, three wells were acquired per tested condition; in each well, nine fields of view were imaged.

**Table S2: Primary and secondary antibodies used in ICC experiments.**

| Primary Antibody | Name | Species | Epitope | Dilution (ICC) |
|---|---|---|---|---|
| | Anti-aSyn total (clone: SYN-1) | Mouse | 91-99 | 1 : 500 |
| | anti-pS129-aSyn (clone: 81A) | Rabbit | pS129 | 1 : 1000 |
| | anti-pS129-aSyn (clone: 81A) | Mouse | AYEMPpSEEGYQ | 1:1000 |
| | anti-NeuN | Rabbit | Synthetic peptide within Human NeuN aa 1-100 | 1 :2000 |
| | anti-MAP2 | Chicken | Recombinant full length | 1 :1000 |
| Secondary Antibody | Goat anti-rabbit Alexa Flour 488 | - | - | 1 : 10000 |
| | Donkey anti-mouse Alexa Fluor 647 | - | - | 1 : 1000 |
| | DAPI | - | - | 1 : 500 |

| | | | | |
|---|---|---|---|---|
| | Phalloidine | - | - | 1 : 500 |
| | Donkey anti-chicken Alexa Fluor 488 | - | - | 1:400 |
| | Donkey anti-rabbit Alexa Fluor 568 | - | - | 1:800 |
| | Goat anti-mouse Alexa Fluor 647 | - | - | 1:800 |

#### 17. UPLC and MS analysis of aSyn double and triple Cys-mutants.

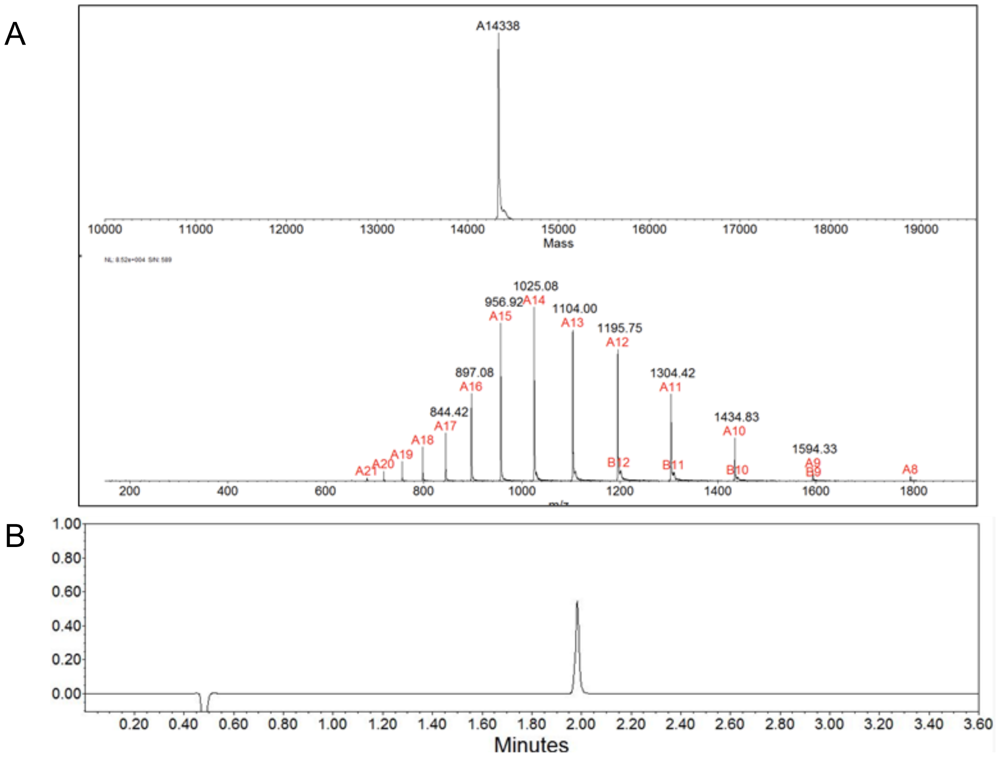


**Figure S5.** A. MS spectrum of purified aSynY125CY133C. Calculated mass = 14340 Da, observed mass = 14338 Da. B. Reversed-phase UHPLC.

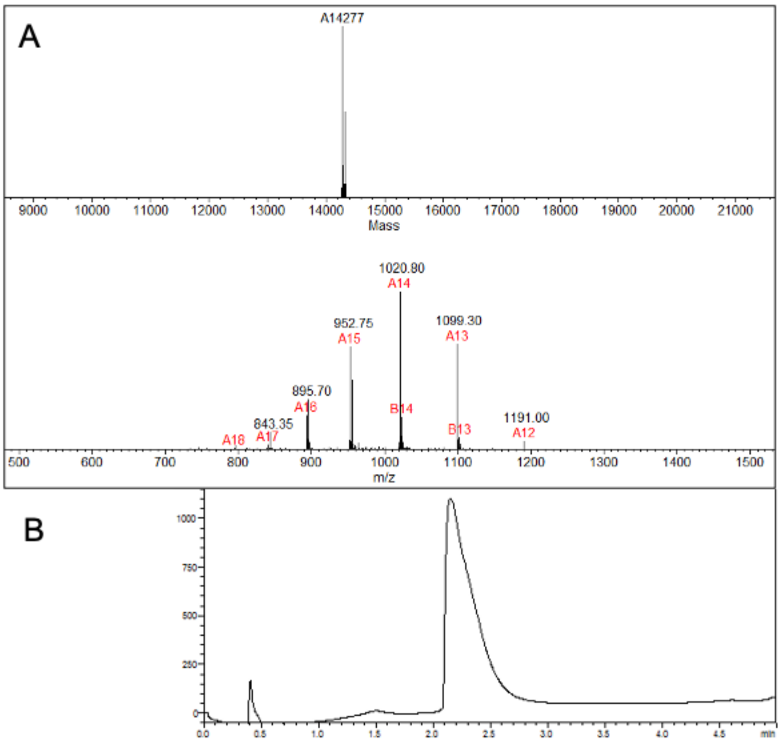


**Figure S6.** A. MS spectrum of purified aSynY125C/Y133CY136C. Calculated mass = 14280 Da, observed mass = 14277 Da. B. Reversed-phase UHPLC.

#### 18. Site-specific aSyn phosphorylation via Pd-OAC-NP

##### 18.1 aSynY39C-P(nat)

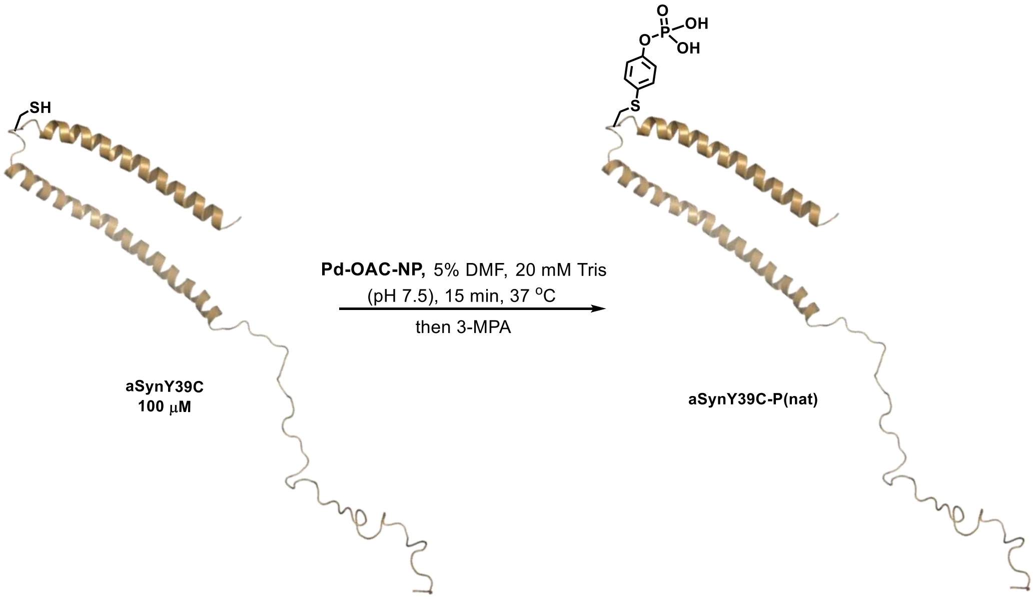


To a 15 mL Falcon tube, aSynY39C (4.3 mg, 2.8 mL, 106 μM, 1.0 equiv.) in 20 mM Tris (pH 7.5) was added, and the aSynY39C solution was preheated at 37 °C for 1 min. Pd-OAC-NP (2.3 mg, 149 μL, 14.8 mM, 7.5 equiv.) as a solution in DMF (v = 5 %) was added in one portion. The final reaction concentrations of the major reaction components were as follows: aSynY39C (100 µM); Pd-OAC-NP (750 µM). The reaction mixture was vigorously stirred for a few seconds and then incubated at 37 °C for 15 min. Then, 3-MPA (0.4 µL, 15 equiv. compared with aSynY39C) was added to quench the reaction, followed by a 15 min incubation at 37 °C. The reaction mixture was purified by RP-HPLC, affording the desired aSynY39C-P(nat) product, 3.5 mg (81%).

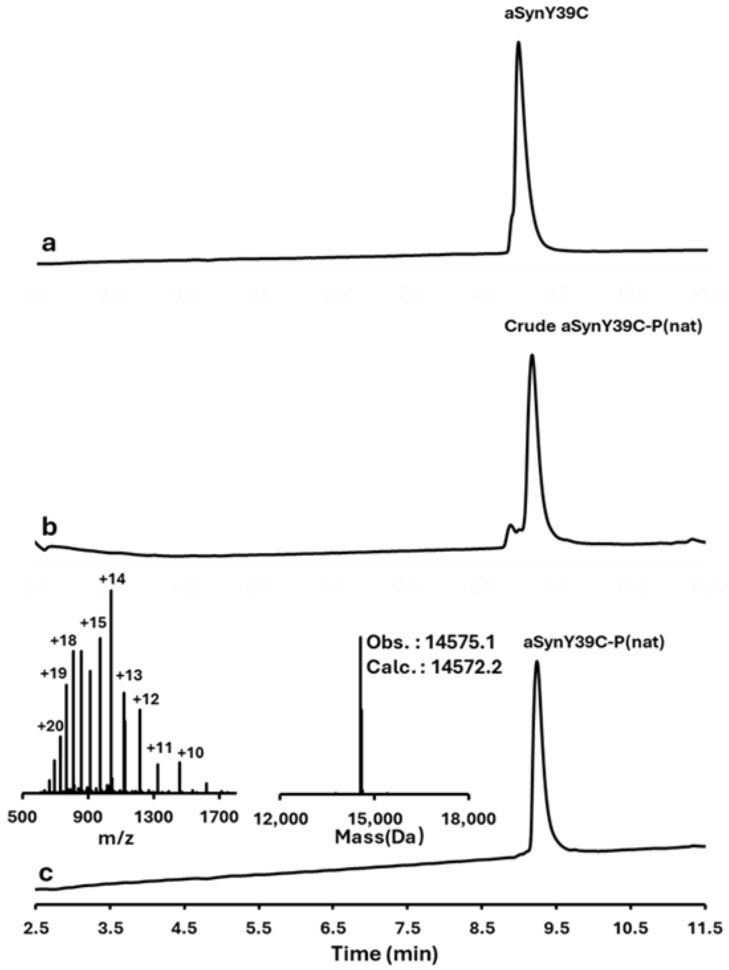


**Figure S7. LCMS analysis of aSynY39C phosphorylation via Pd-OAC-NP.** LC chromatogram of the UV absorbance at 214 nm, mass-to-charge (m/z) spectrum, and deconvoluted mass spectrum. (a) aSynY39C at t = 0. (b) crude native phosphorylation reaction of aSynY39C with Pd-OAC-NP at t = 15 min. (c) isolated aSynY39C-P(nat).

##### 18.2 aSynY125C-P(nat)

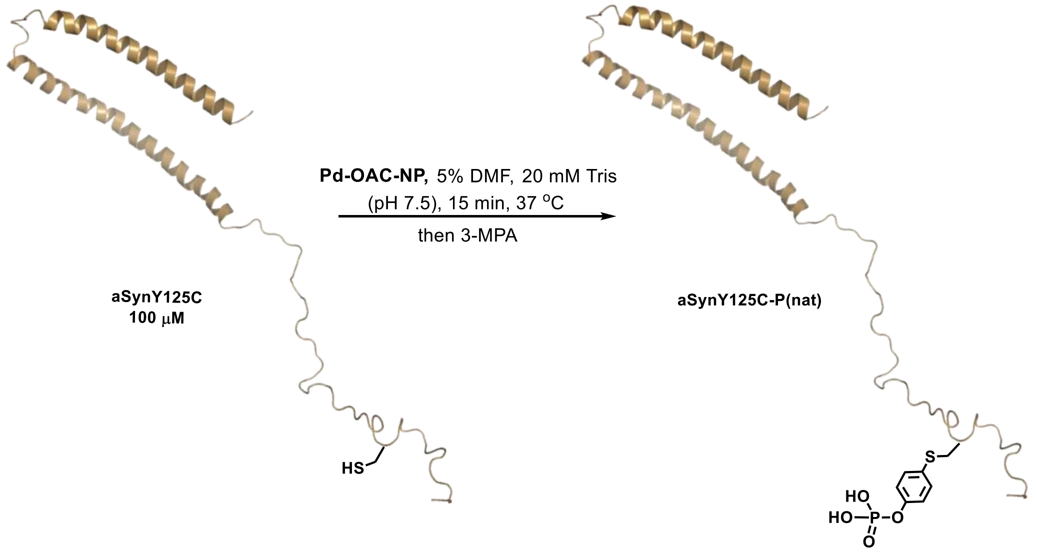


To a 15 mL Falcon tube, aSynY125C (4.2 mg, 2.7 mL, 105 μM, 1.0 equiv.) in 20 mM Tris (pH 7.5) was added, and the solution was preheated at 37 °C for 1 min. Then Pd-OAC-NP (2.1 mg, 145 μL, 14.7 mM, 7.5 equiv.) as a solution in DMF (v = 5 %) was added in one portion. The final reaction concentrations of the major reaction components were as follows: aSynY125C (100 μM); Pd-OAC-NP (750 μM). The reaction mixture was vigorously stirred and followed by an incubation at 37 °C for 15 min. Then, 3-MPA (0.4 μL, 15 equiv. compared withaSynY125C) was added, and the reaction mixture was kept at 37 °C for 15 min to quench the reaction. The reaction mixture was purified by RP-HPLC using method B, affording the desired product aSynY125C-P(nat), 3.7 mg (89%).

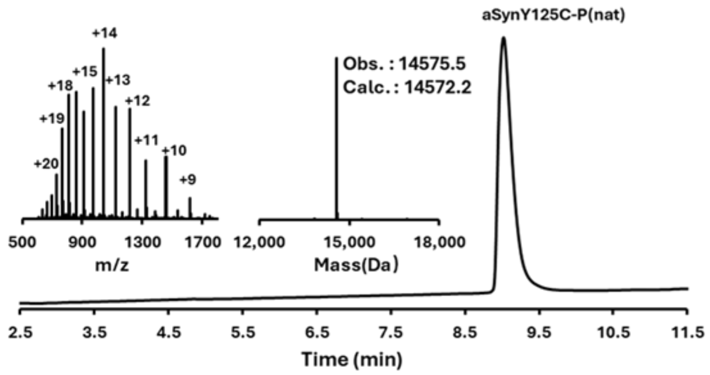


**Figure S8. LCMS analysis of aSynY125C phosphorylation via Pd-OAC-NP.** LC chromatogram of the UV absorbance at 214 nm, mass-to-charge (m/z) spectrum, and deconvoluted mass spectrum.

##### 18.3 aSynY133C-P(nat)

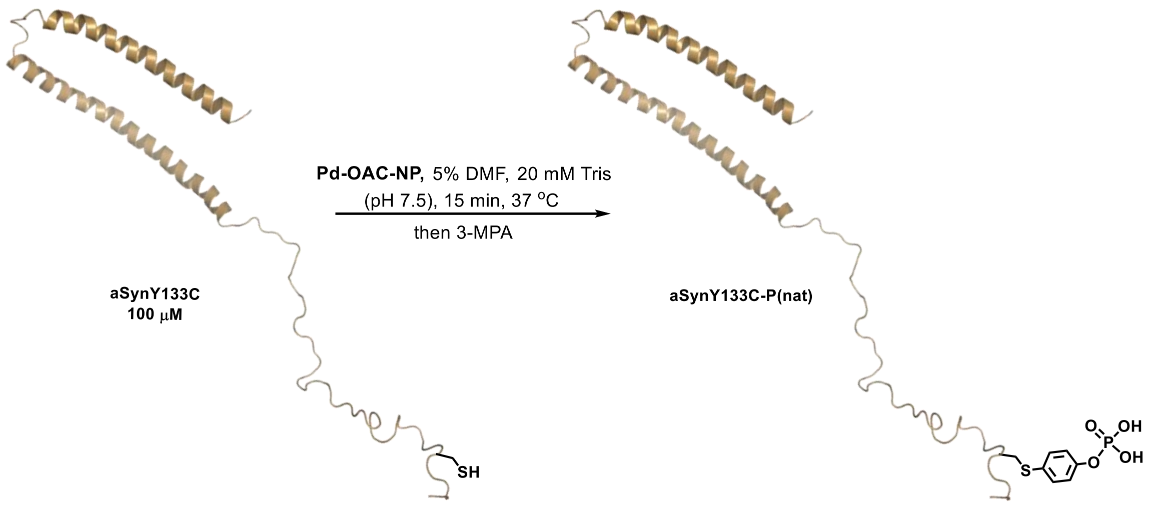


To a 15 mL Falcon tube, **aSynY133C** (4.0 mg, 2.6 mL, 105 µM, 1.0 equiv.) in 20 mM Tris (pH 7.5) was added, and the solution was preheated at 37 °C for 1 min. Then **Pd-OAC-NP** (2.1 mg, 139 µL, 14.8mM, 7.5 equiv.) as a solution in DMF (v = 5 %) was added in one portion. The final reaction concentrations of the major reaction components were as follows: **aSynY133C** (100 µM); **Pd-OAC-NP** (750 µM). The reaction mixture was vigorously stirred for a few seconds and then kept at 37 °C for 15 min. Then, 3-MPA (0.4 µL, 15 equiv. compared with **aSynY133C)** was added, and the reaction mixture was kept at 37 °C for 15 min to quench the reaction. The reaction mixture was purified by RP-HPLC using method B, affording the desired product **aSynY133C-P(nat),** 3.5 mg (87%).

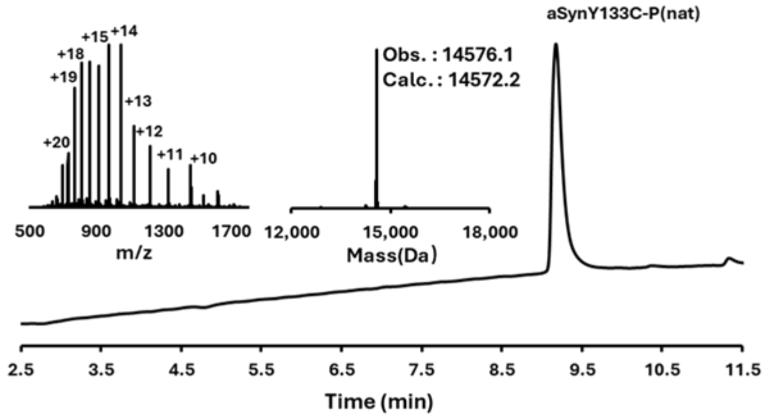


**Figure S9. HPLC and MS analysis of aSynY133C phosphorylation via Pd-OAC-NP.** LC chromatogram of the UV absorbance at 214 nm, mass-to-charge (m/z) spectrum, and deconvoluted mass spectrum.

#### 18.4 aSynY136C-P(nat)

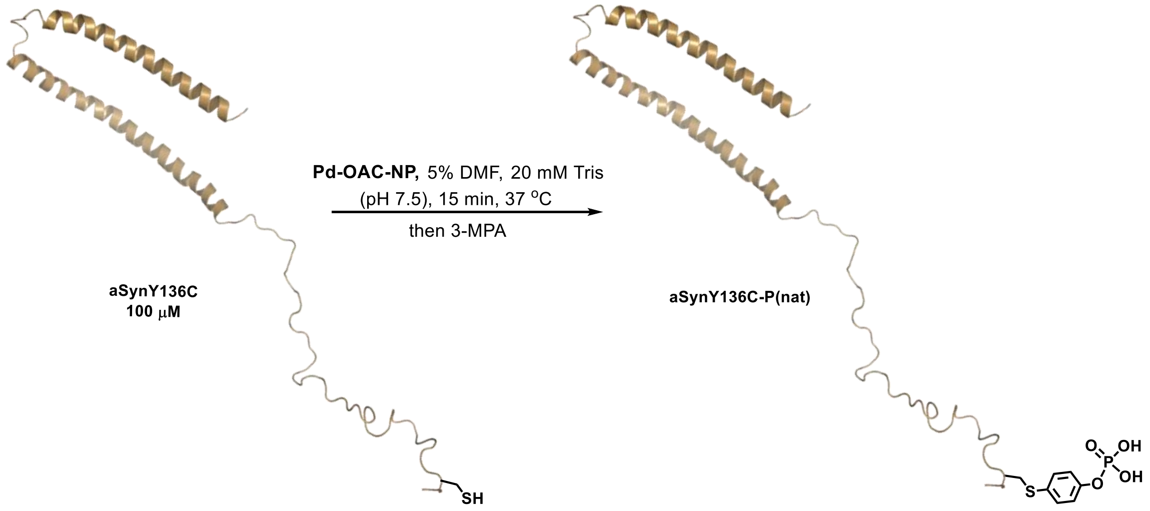


To a 15 mL Falcon tube, aSynY136C (4.0 mg, 2.6 mL, 105.1 µM, 1.0 equiv.) in 20 mM Tris (pH 7.5) was added, and the solution was preheated at 37 °C for 1 min. Then, Pd-OAC-NP (2.1 mg, 139 µL, 14.8 mM, 7.5 equiv.) as a solution in DMF (v = 5 %) was added in one portion. The reaction mixture was vigorously stirred for a few seconds and then kept at 37 °C for 15 min. The final reaction concentrations of the major reaction components were as follows: aSynY136C (100 µM); Pd-OAC-NP (750 µM). Then, 3-MPA (0.4 µL, 15 equiv. compared with aSynY136C) was added, and the reaction mixture was kept at 37 °C for 15 min to quench the reaction. The reaction mixture was purified by RP-HPLC using method B, which afforded the desired product aSynY136C-P(nat) 3.3 mg (82%).

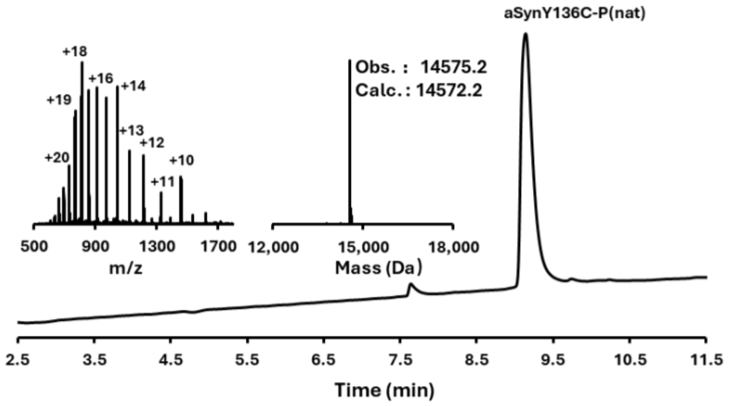


**Figure S10. HPLC and MS analysis of aSynY136C phosphorylation via Pd-OAC-NP.** LC chromatogram of the UV absorbance at 214 nm, mass-to-charge (m/z) spectrum, and deconvoluted mass spectrum.

#### 18.5 aSynY125CY133C-P(nat) [di-phosphorylation]

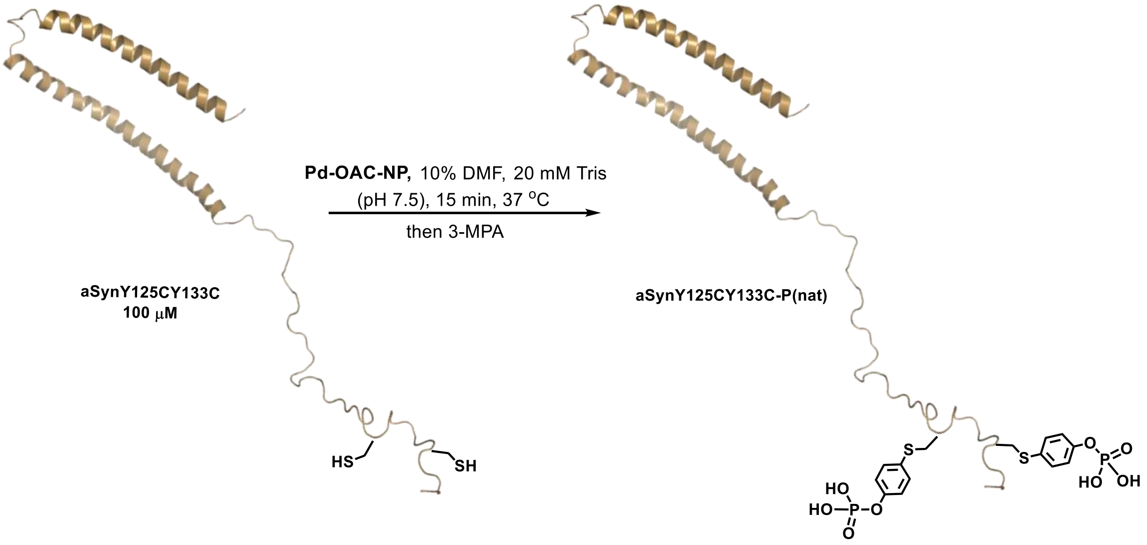


To a 15 mL Falcon tube, aSynY125CY133C (2.0 mg, 1.25 mL, 111.1 μM, 1.0 equiv.) in 20 mM Tris (pH 7.5) was added, and the solution was preheated at 37 °C for 1 min. Then, Pd-OAC-NP ( 2.9 mg, 139 μL, 20 mM, 20 equiv.) as a solution in DMF (v = 10 %) was added in one portion. The reaction mixture was vigorously stirred for a few seconds, then incubated at 37 °C for 20 min. The final reaction concentrations of the major reaction components were as follows: aSynY125CY133C (100 μM); Pd-OAC-NP (2 mM). Then, 3-MPA (0.4 μL, 30 equiv. relative to aSynY125CY133C) was added, and the reaction mixture was incubated at 37 °C for 20 min to quench the reaction. The reaction mixture was purified by RP-HPLC using method B, affording the desired product aSynY125CY133C-P(nat), 0.9 mg (45 %).

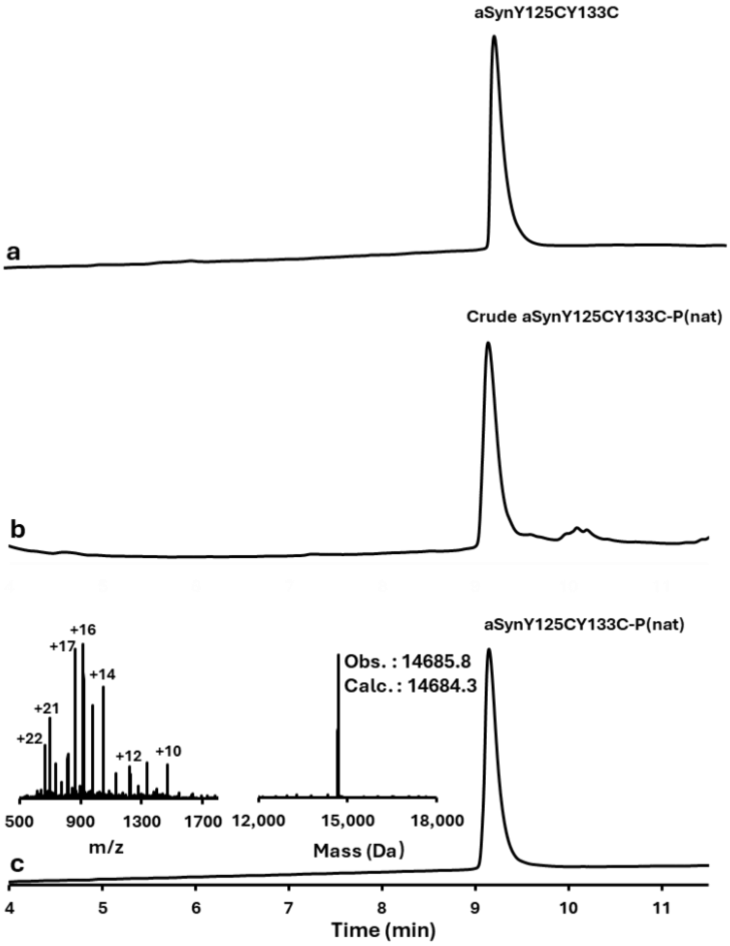


**Figure S11. LCMS analysis of aSynY125CY133C phosphorylation via Pd-OAC-NP.** LC chromatogram of the UV absorbance at 214 nm, mass-to-charge (m/z) spectrum, and deconvoluted mass spectrum. (a) aSynY125CY133C at t = 0. (b) crude dual native phosphorylation reaction of aSynY125CY133C with Pd-OAC-NP at t = 15 min. (c) isolated aSynY125CY133C-P(nat).

#### 18.6 aSynY125CY133CY136C-P(nat) [tri-phosphorylatio]

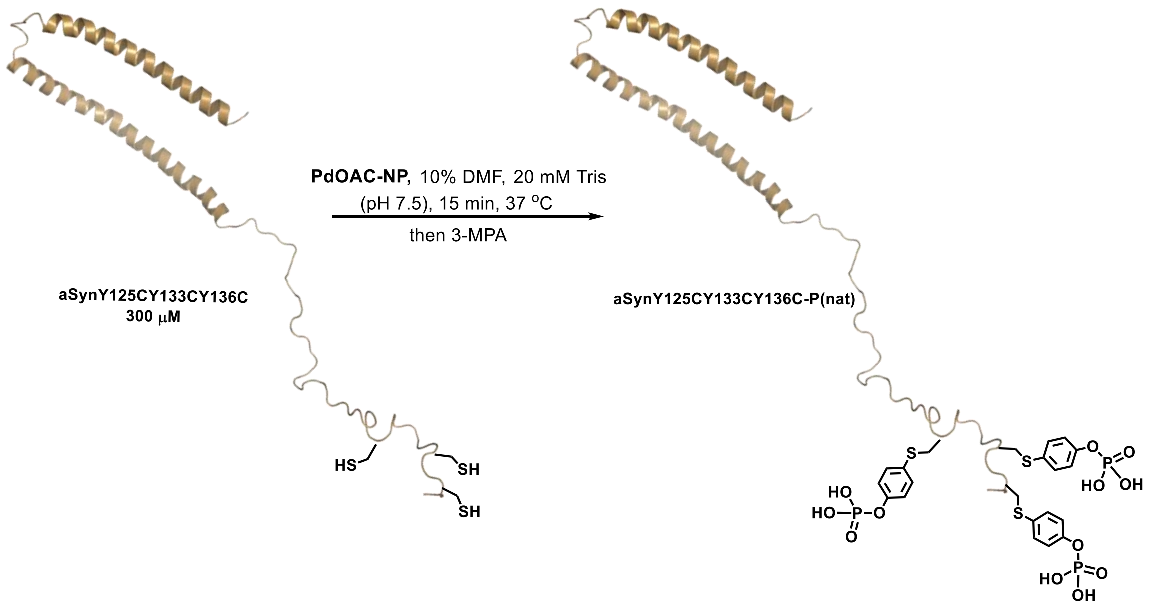


To a 15 mL Falcon tube, aSynY125CY133CY136C (3.8 mg, 798 μL, 333 μM, 1.0 equiv.) in 20 mM Tris (pH 7.5) was added, and the solution was preheated at 37 °C for 1 min. Then Pd-OAC-NP (6.2 mg, 89 μL, 67 mM, 22.5 equiv.) as a solution in DMF (v = 10 %) was added in one portion. The reaction mixture was vigorously stirred for a few seconds and then kept at 37 °C for 15 min. The final reaction concentrations of the major reaction components were as follows: aSynY136C (300 μM); Pd-OAC-NP (6.75 mM). Then, 3-MPA (0.53 μL, 22.5 equiv. compared with aSynY125C/Y133C/Y136C) was added, and the reaction mixture was kept at 37 °C for 15 min to quench the reaction. The reaction mixture was purified by RP-HPLC using method B, affording the desired product aSynY125CY133CY136C-P(nat) 2.1 mg (54%).

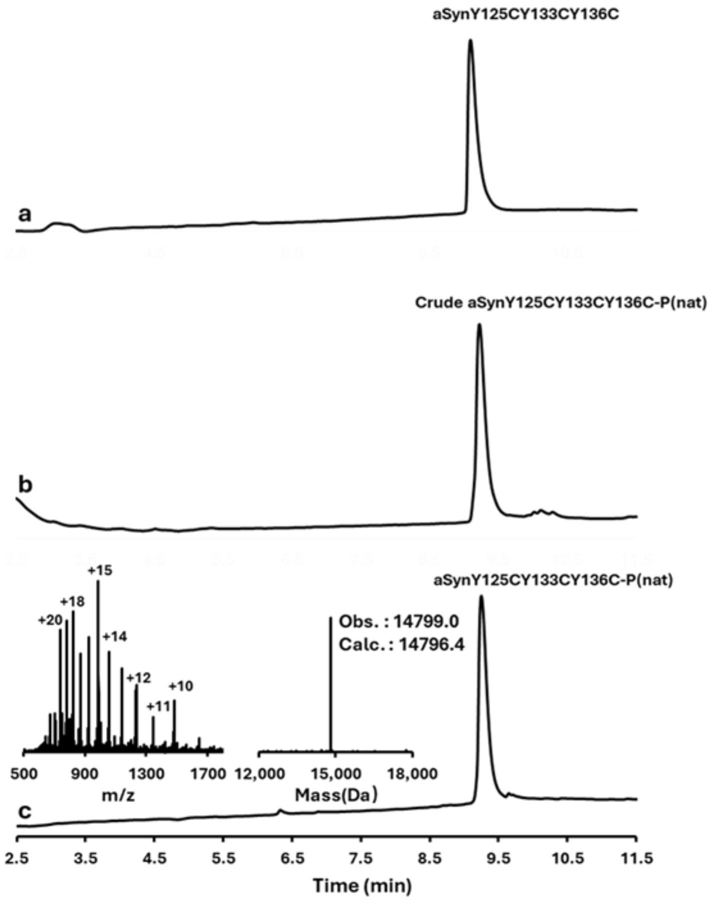


**Figure S12. LCMS analysis of aSynY125CY133CY136C phosphorylation via Pd-OAC-NP.** LC chromatogram of the UV absorbance at 214 nm, mass-to-charge (m/z) spectrum, and deconvoluted mass spectrum. (a) aSynY125CY133CY136C at t = 0. (b) crude triple phosphorylation reaction of aSynY125CY133CY136C with Pd-OAC-NP at t = 15 min. (c) isolated triple phosphorylated aSynY125CY133CY136C-P(nat).

#### 19. SDS-PAGE analysis of different aSyn analogues: WT, mono-P(nat), di-P(nat), and tri-P(nat)

A.

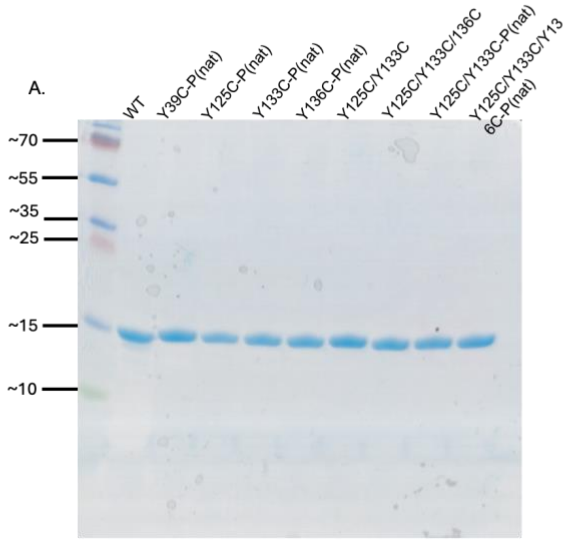


B.

| Protein | Molecular weight (observed) | Molecular weight (calculated) |
|---|---|---|
| h-aSyn-WT | 14460 Da | 14460 Da |
| h-aSyn-Y39C-P(nat) | 14575 Da | 14572 Da |
| h-aSyn-Y125C-P(nat) | 14575 Da | 14572 Da |
| h-aSyn-Y133C-P(nat) | 14576 Da | 14572 Da |
| h-aSyn-Y136C-P(nat) | 14575 Da | 14572 Da |
| h-aSyn-Y125C/Y133C | 14338 Da | 14340 Da |
| h-aSyn-Y125/-Y133C/Y136C | 14'279 Da | 14'280 Da |
| h-aSyn-Y125C/Y133C-P(nat) | 14585 Da | 14684 Da |
| h-aSyn-Y125/-Y133C/Y136C-P(nat) | 14799 Da | 14796 Da |

**Figure S13.** (A) Gel electrophoresis of WT, mono-P(nat), di-Cys mutant, and P(nat), tri-Cys mutant, and P(nat) analogues. The compounds were loaded onto a 16% tricine gel and run for 2 hours at 120 V. (B) Molecular weight of the corresponding protein.

#### 20. Dot blot to validate the detection of aSyn-P(nat) via antibodies and phosphatase activity:

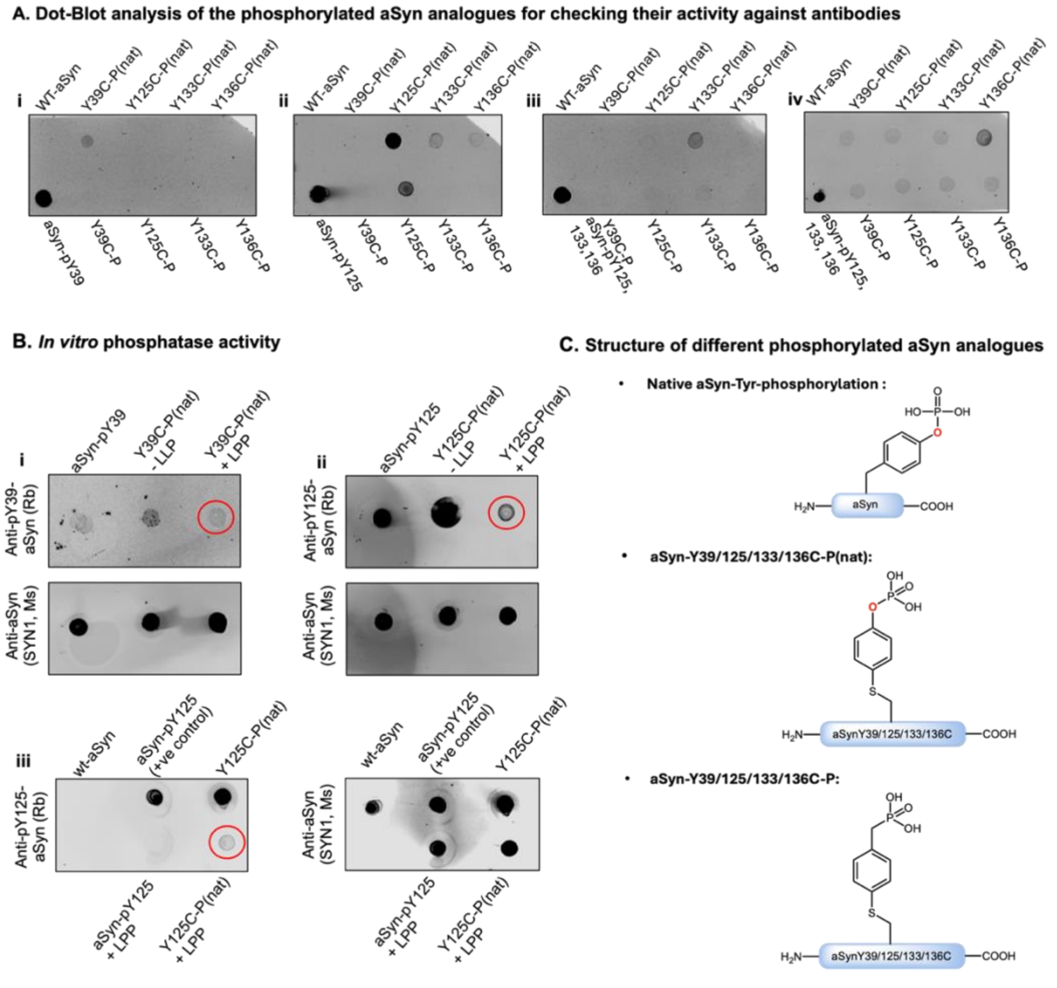


**Figure S14. The activity of the synthetic phosphorylated aSyn analogues.** (A) Dot-blot analysis of the phosphorylated *a*Syn(h) analogues (20 µM each) against (i) anti-Syn-pY39, (ii) anti-Syn-pY125, (iii) anti-Syn-pY133, (iv) anti-Syn-pY133, Syn-pY136. (B) In vitro phosphatase activity of (i) aSyn-Y39C-P(nat) and (ii, iii) aSyn-Y125C-P(nat) in the presence of Lambda Protein Phosphatase (1x-LLP). In Biii, 3x LLP concentration was used to get full dephosphorylation. (C) Structure of native Tyr-phosphorylated aSyn, aSyn-Y39/125/133/136C-P(nat), and aSyn-Y39/125/133/136C-P.

#### 21. Aggregation ThT assay [WT vs. mono-P(nat) analogues]:

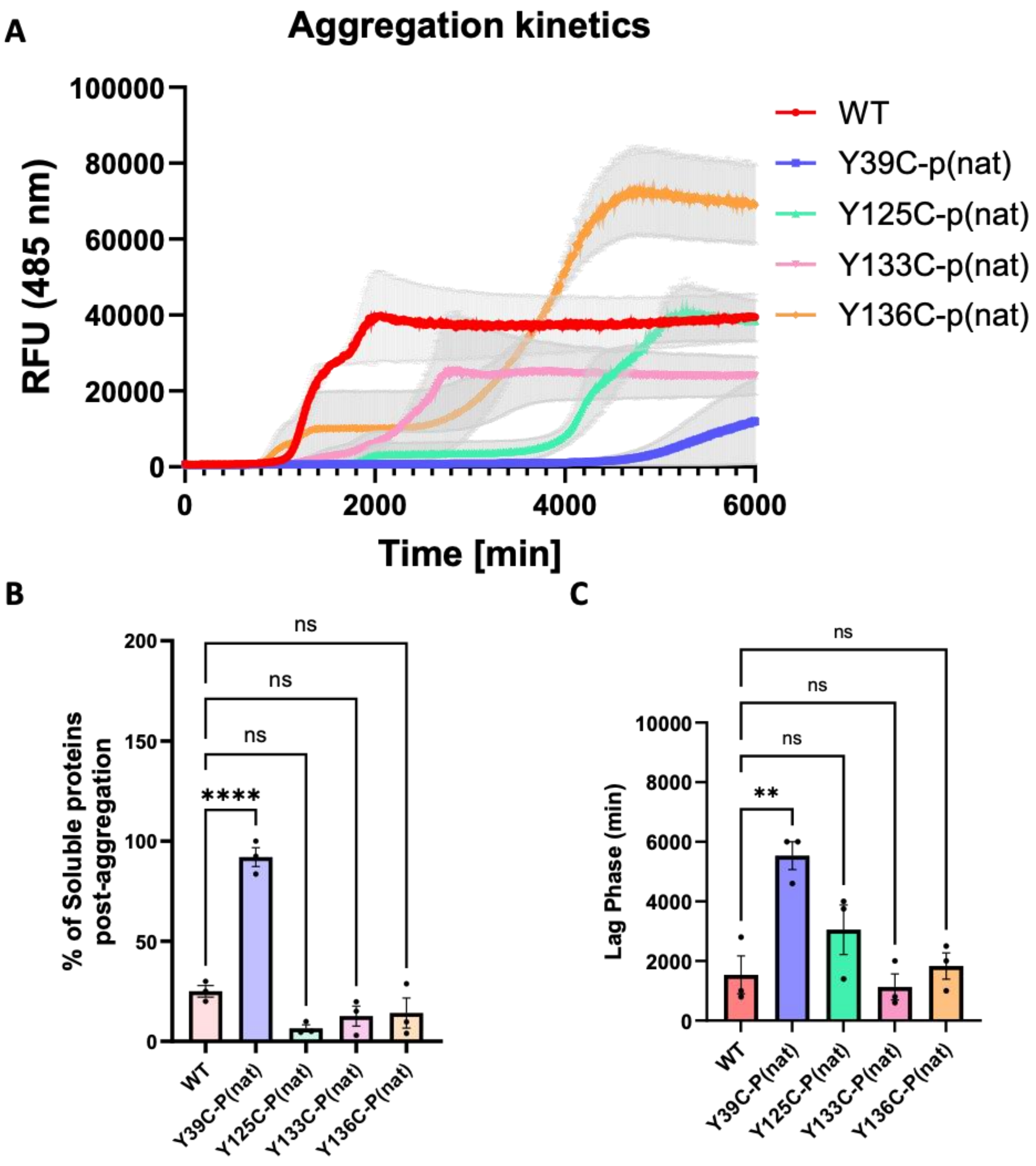


**Figure S15.** (A) Aggregation kinetics of WT and mono-phosphorylated aSyn [Y39C-P(nat), Y125C-P(nat), Y133C-P(nat), Y136C-P(nat)] at 20 µM initial concentration monitored by ThT fluorescence. RFU, relative fluorescence units, n = 3. (B) Solubility assay showing the percentage of the soluble protein post-aggregation of all the aSyn analogues (at the final time point). (C) Bar graph of lag phase extracted from the ThT aggregation kinetics shown by (A) (mean ± SEM, n = 3).

#### 22. Sedimentation Assay for aSyn WT, mono-P(nat), di-P(nat) and tri-P(nat) post-fibrilization:

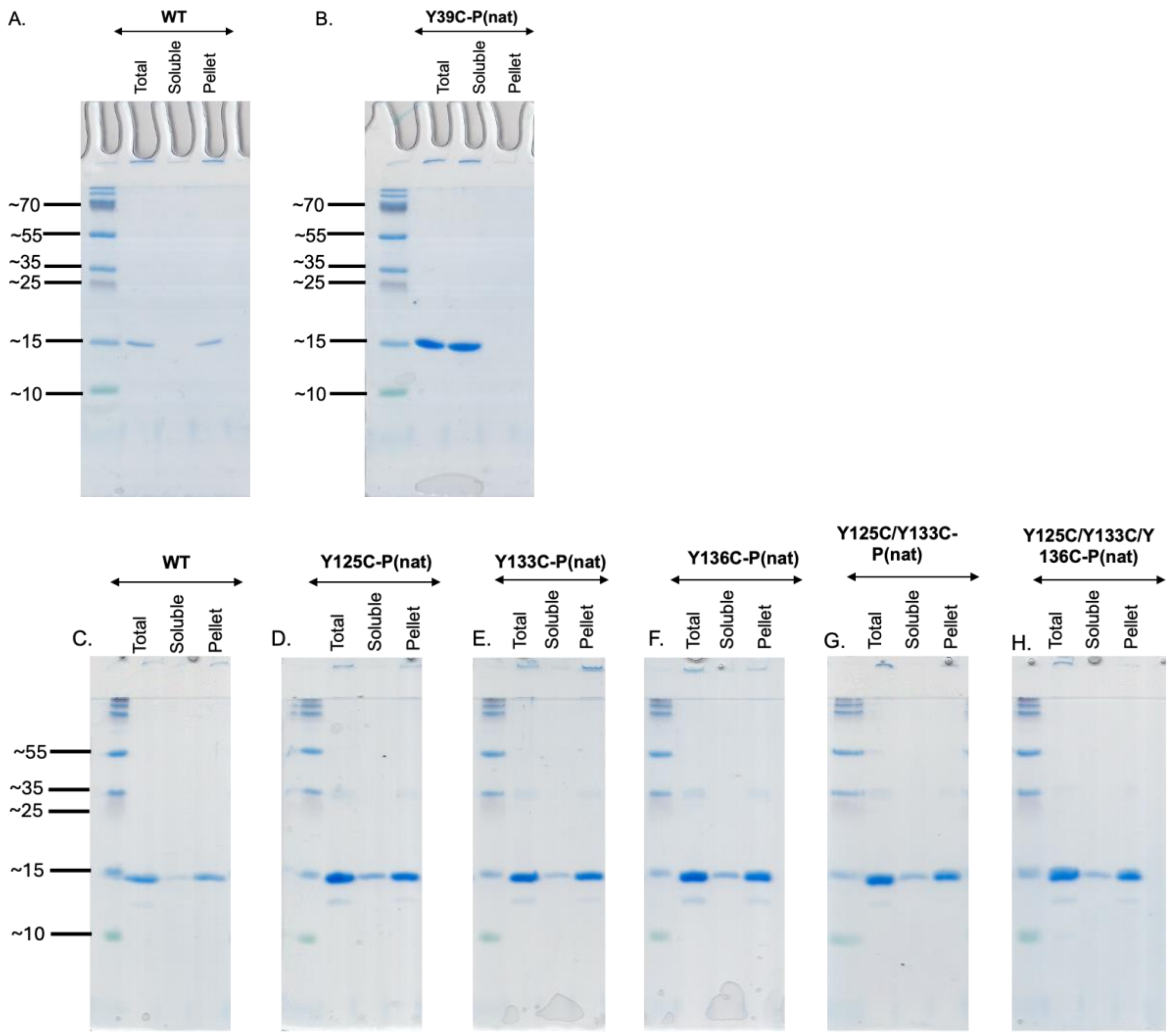


**Figure S16.** Separation of fibrils from the soluble (monomer and oligomer) species for aSyn. (A, C) WT, (B) Y39C-P(nat), (D) Y125C-P(nat), (E) Y133C-P(nat), (F) Y136C-P(nat), (G) Y125C/Y133C-P(nat), (H) Y125C/Y133C/136C-P(nat).

#### 23. Characterization of the fibrils of aSyn WT, mono-P(nat), di-P(nat) and tri-P(nat) post-aggregation:

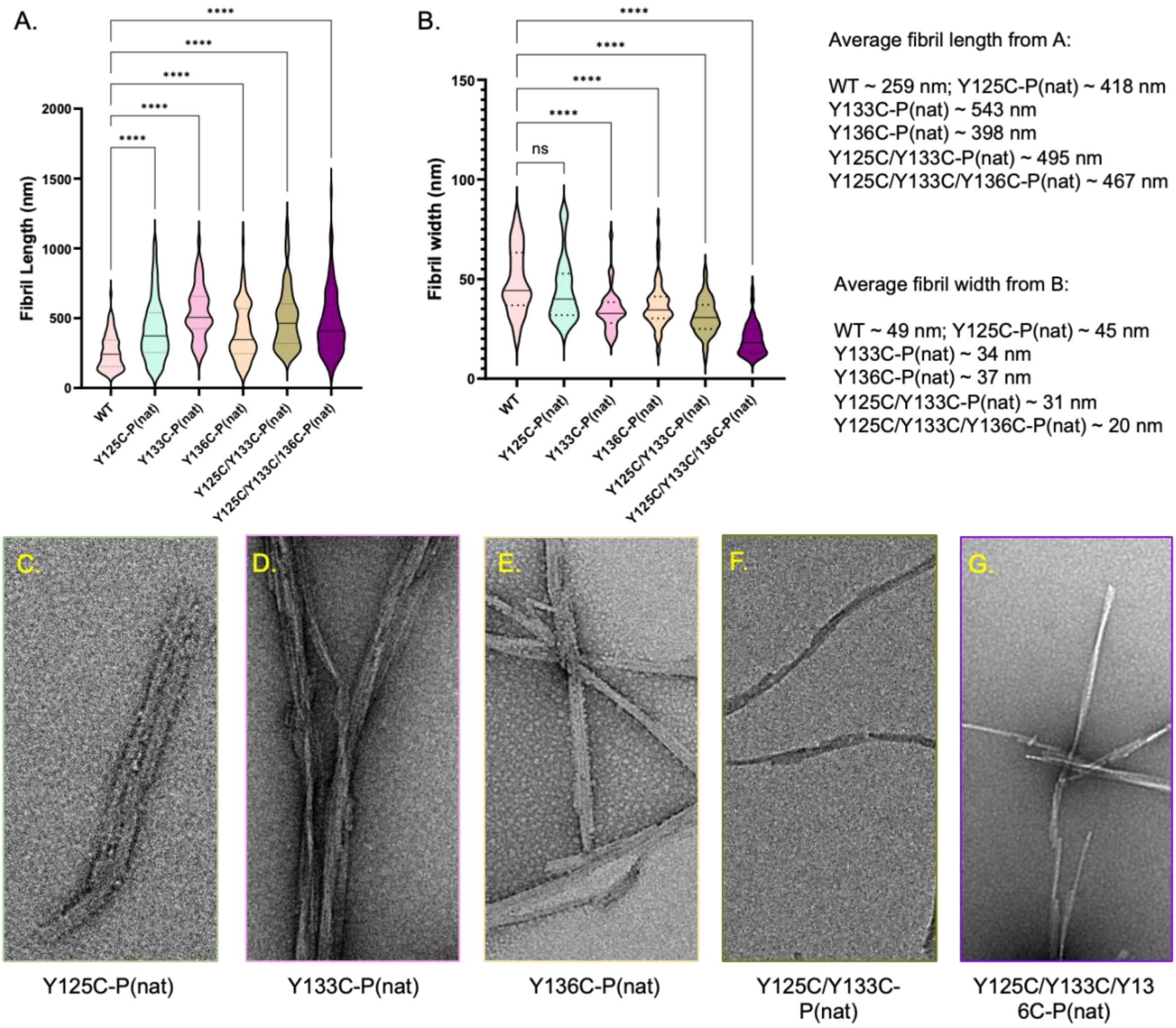


**Figure S17. Characterization of the fibrils**. (A) Violin plot of the length of the aSynWT, Y125C-P(nat), Y133C-P(nat), Y136C-P(nat), Y125C/Y133C-P(nat), Y125C/Y133C/Y136C-P(nat). (B) Violin plot of the length of the aSynWT, Y125C-P(nat), Y133C-P(nat), Y136C-P(nat), Y125C/Y133C-P(nat), Y125C/Y133C/Y136C-P(nat). (C-G) Zoomed images of the fibrils of Y125C-P(nat), Y133C-P(nat), Y136C-P(nat), Y125C/Y133C-P(nat), Y125C/Y133C/Y136C-P(nat), respectively.

#### 24. Purity and characterization of aSyn PFFs:

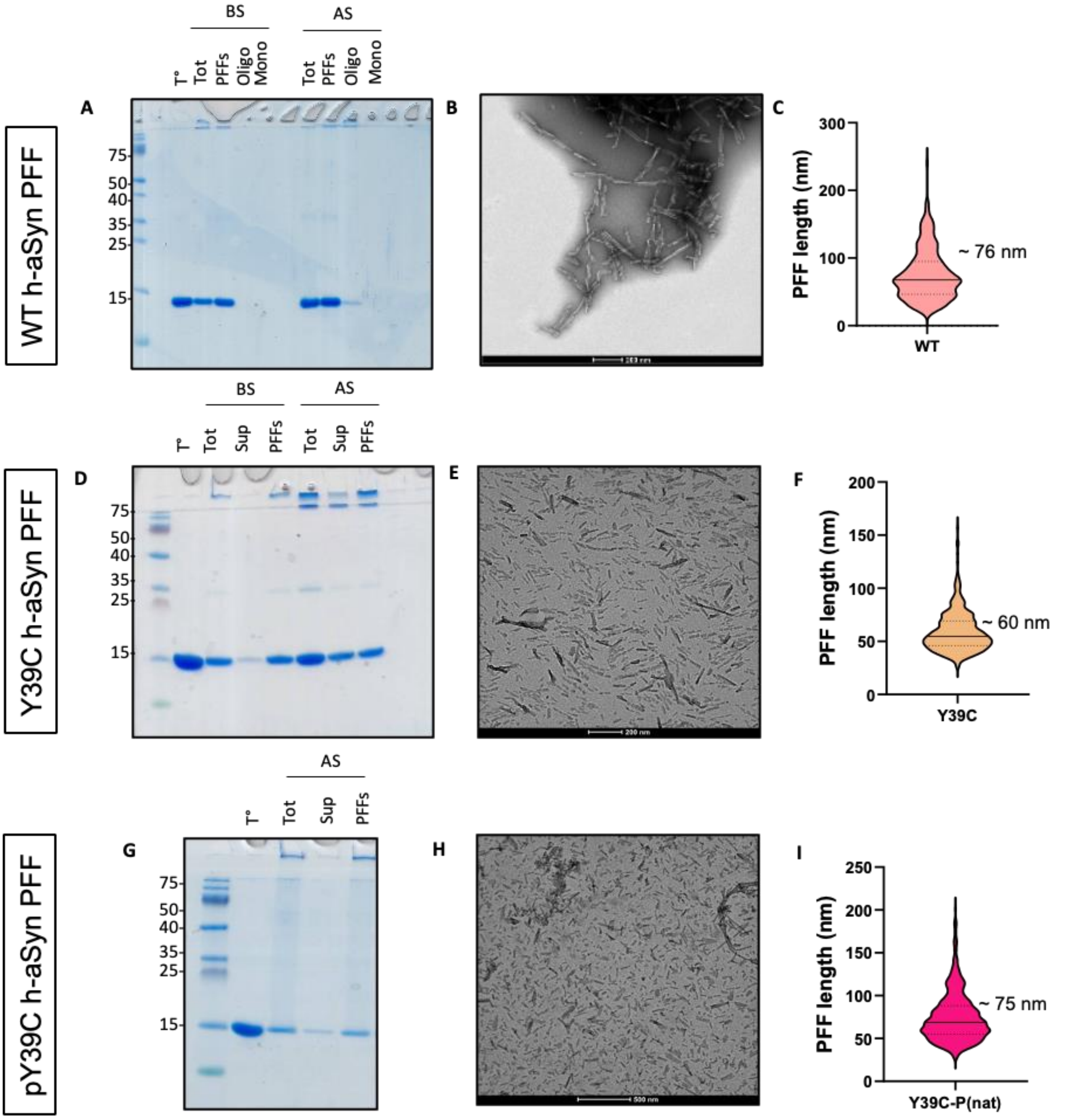


**Figure S18.** aSyn WT, Y39C, and Y39C-P(nat) fibrils were formed by incubation of monomeric aSyn for 5 days at 37°C under constant agitation at 1000 rpm. **A.** Purity of α-syn fibrils was verified by SDS-PAGE and Coomassie blue staining. After sonication, fibril preparations were centrifuged, and the presence of the fibrils was verified in the pellet fraction, while the absence of monomer release after the sonication step was assessed in the supernatant fraction or after filtration through a 100 kDa filter (filtration). **B.** aSyn fibrils were characterized by transmission electron microscopy (TEM) imaging. Representative images of negatively stained aSyn fibrils after sonication. All aSyn fibrils showed the characteristic rigid, non-branched fibrillar morphology. Scale bars = 200 nm. **C.** Average length of the fibrils after sonication. BS = before sonication, AS = after sonication.

#### 25. Generation of WT aSyn (h) stably expressing U2OS cell line:

For the expression, the HTB-96 U2OS Osteosarcoma Human cell line from ATCC, with lot number 70025046, was used. Lentivirus (686-pLenti-aSyn-WT) was produced in the lab, virus titrated, and viral DNA measured by qPCR showed 5.00E+06 vg/mL viral genome in the lentivirus. For infection, 686-pLenti-aSyn-WT 5.00E+06 vg/mL lentivirus carrying the selection marker for puromycin was used at a Multiplicity of Infection (MOI) of 1. After infection, cells were allowed to grow and express the protein for puromycin resistance under non-selective conditions for approximately 48 hours, and then treated with puromycin at 8 µg/ml. Cell clones were analysed or further expanded as soon as the cells in the non-transduced control were dead. To generate monoclonal human wild-type alpha-Synuclein stable expressing U2OS cells, two approaches were evaluated. Initial attempts with serial dilution plating directly in 96-well plates did not yield monoclonal populations. Subsequently, single colonies resistant to 8 µg/mL puromycin were isolated after growth in selective media. These colonies were individually picked using trypsin, diluted, and replated in a 96-well format for expansion. Cells were allowed to grow for 4 weeks with bi-weekly replenishment of selective media until 70% confluency was reached, resulting in the successful establishment of monoclonal stable U2OS lines expressing WTaSyn (h).

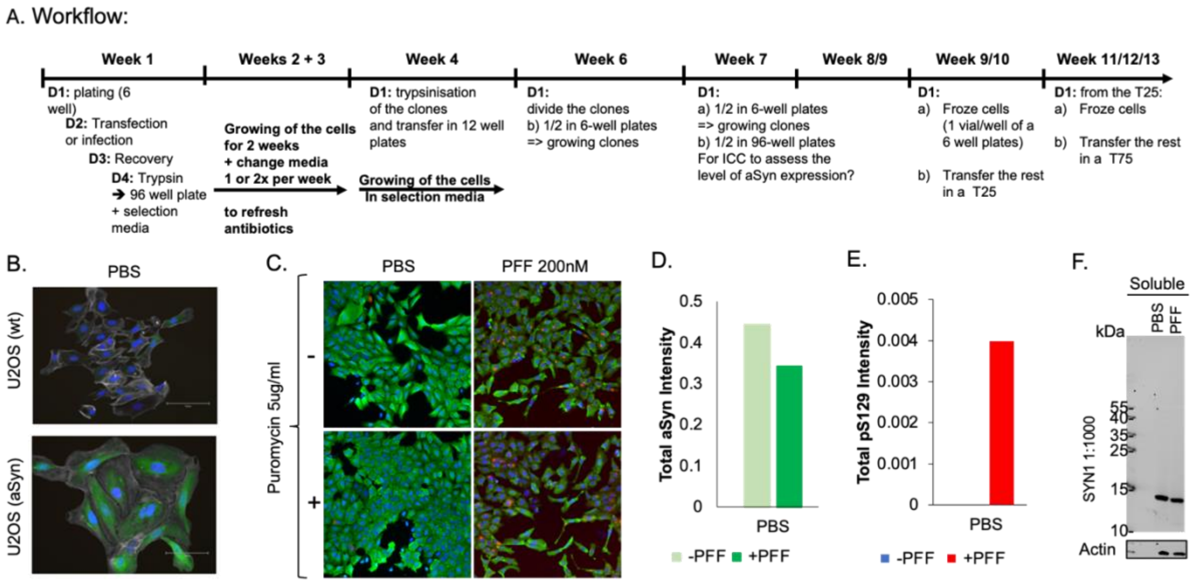


**Figure S19.** (A) Complete workflow for the generation of WT aSyn (h) stable expression in U2OS cells. (B) Validation via ICC of aSyn expression compared to the U2OS (wt) cells (polyclonal). (DAPI: blue, Phalloidin: grey, aSyn: green) (C) Cultured U2OS (aSyn) cells expressing aSyn stably (monoclonal cells), as verified with and without Puromycin (5 µg/mL) addition to the culture media. Our cell line exhibits pS129-positive punctate signals upon treatment with aSyn (wt) PFF (200 nM). (DAPI: blue, pS129: red, aSyn: green). HTS analysis to show (D) the total aSyn expression level and (E) pS129 level with and without the PFF treatment. (F) Western blot against anti-alpha-Synuclein (SYN1) antibody to validate the expression of aSyn in the U2OS (aSyn) cells.

#### 26. Representative images of pS129 pathology in U2OS (aSyn) cells treated with 200 nM PFFs:

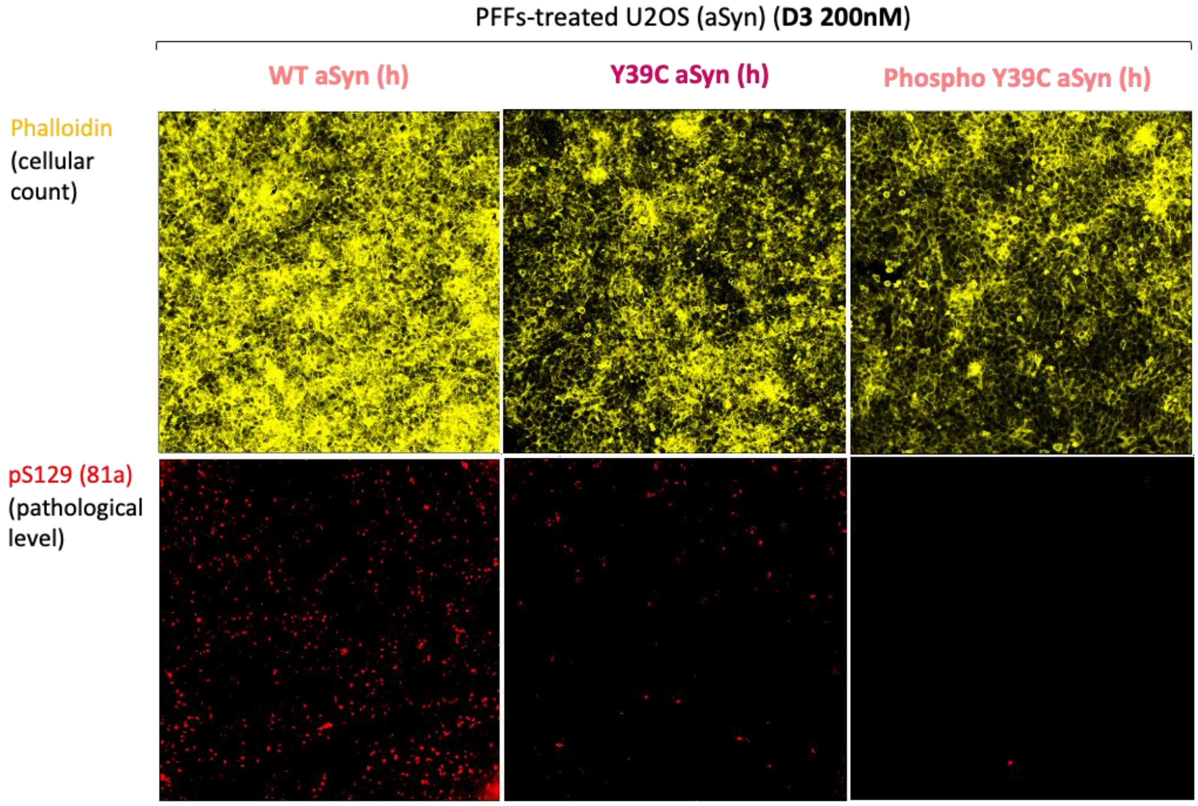


**Figure S20.** Representative images with Phalloidine (yellow) and pS129 (red) staining of the cells treated with WT, Y39C, Y39C-P(nat) PFFs.

#### 27. Representative images of pS129 pathology in U2OS (aSyn) cells treated with 400 nM PFFs:

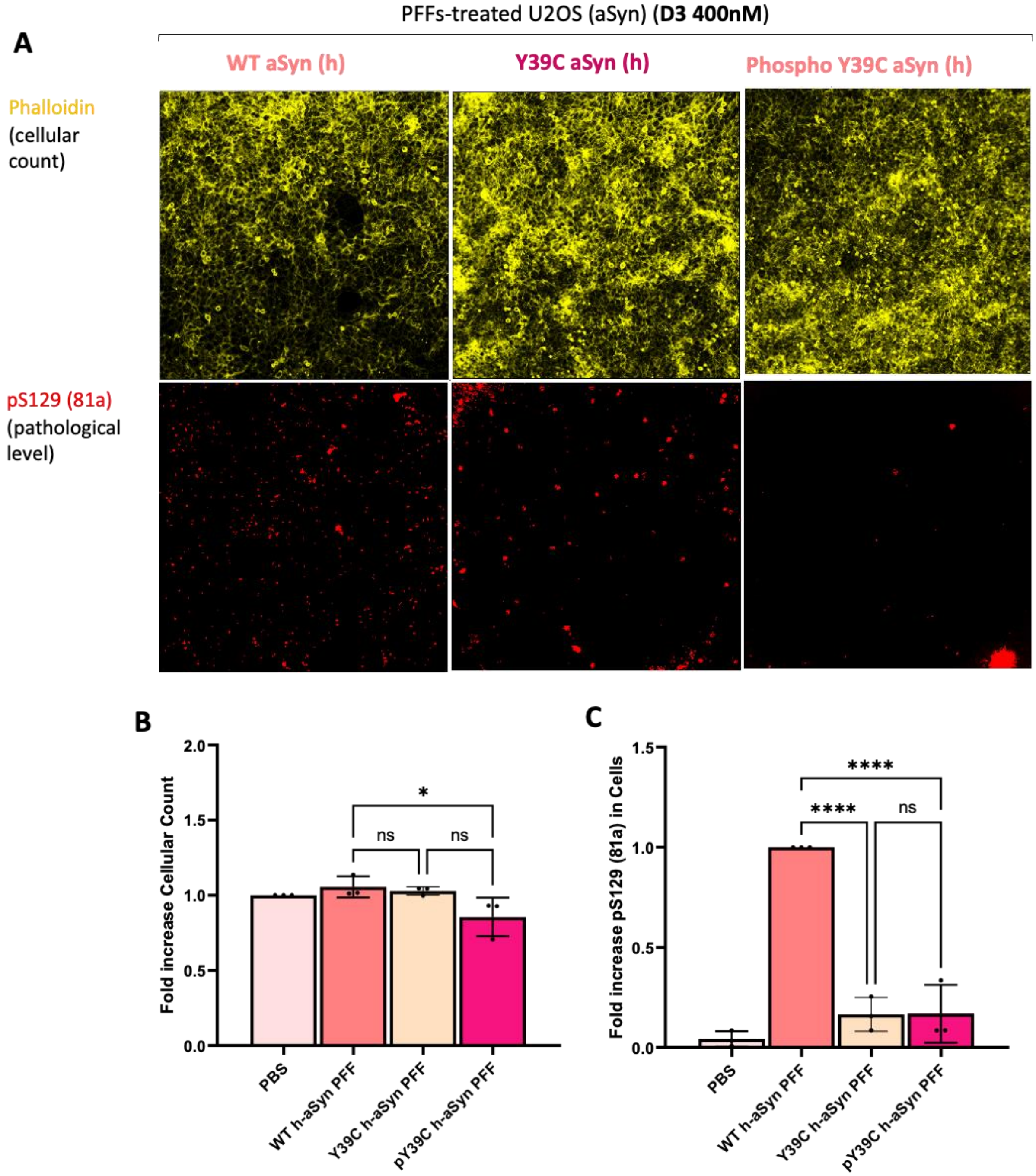


**Figure S21.** Cells are treated with 400nM of either aSyn WT, Y39C, Y39C-P(nat) PFFs or PBS (-ve control). Three days after treatment (D3), the cells are fixed in 4% PFA, and immunocytochemistry is performed. (A) Representative image (Phalloidin = yellow, pS129 = red). (B**)** Fold Increase of cellular count for each condition (Phalloidin), calculated from cells treated with PBS. (one-way ANOVA, $F(3,8) = 4.315$, $P = 0.0436$, $N = 3$). (C**)** Fold Increase of total pS129 level in cells (81a), calculated from cells treated with recombinant human aSyn. (one-way ANOVA, $F(3,8) = 79.40$, $P=<0.0001$, $N = 3$).

#### 28. NMR spectra of synthesized compounds:

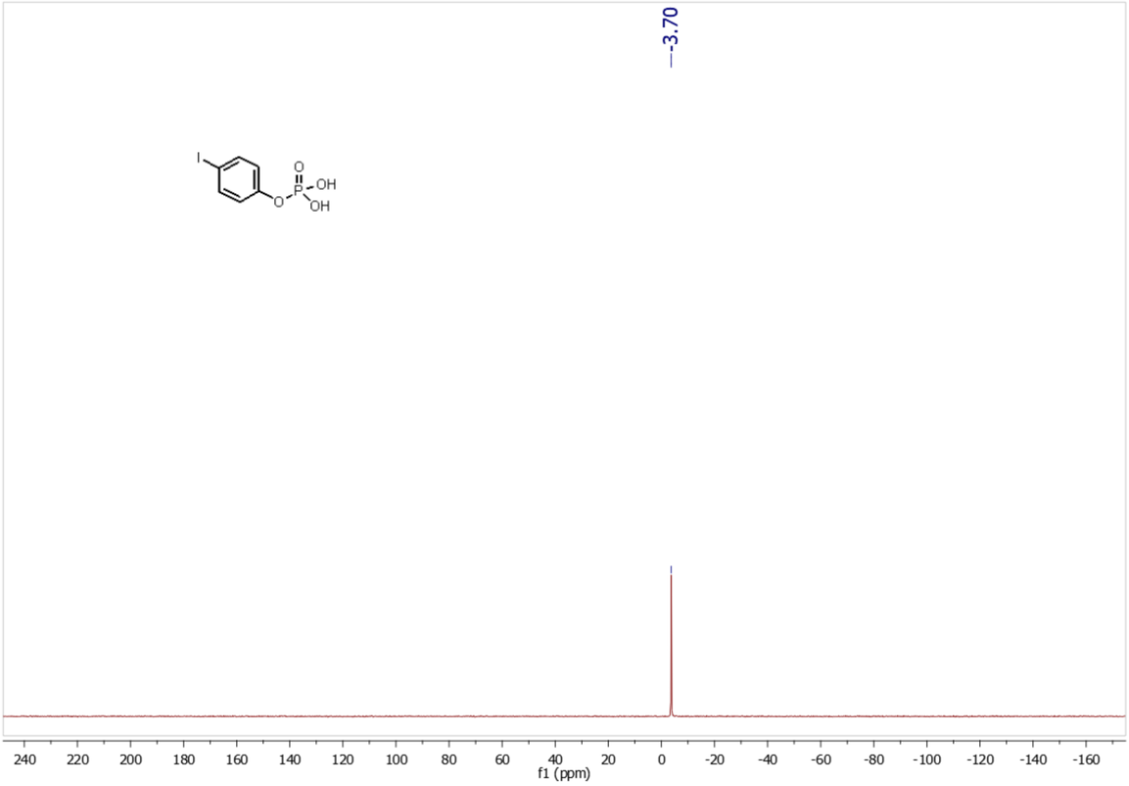


**Figure S22.** $^{31}$P NMR spectrum of 4-iodophenyl dihydrogen phosphate in MeOD.

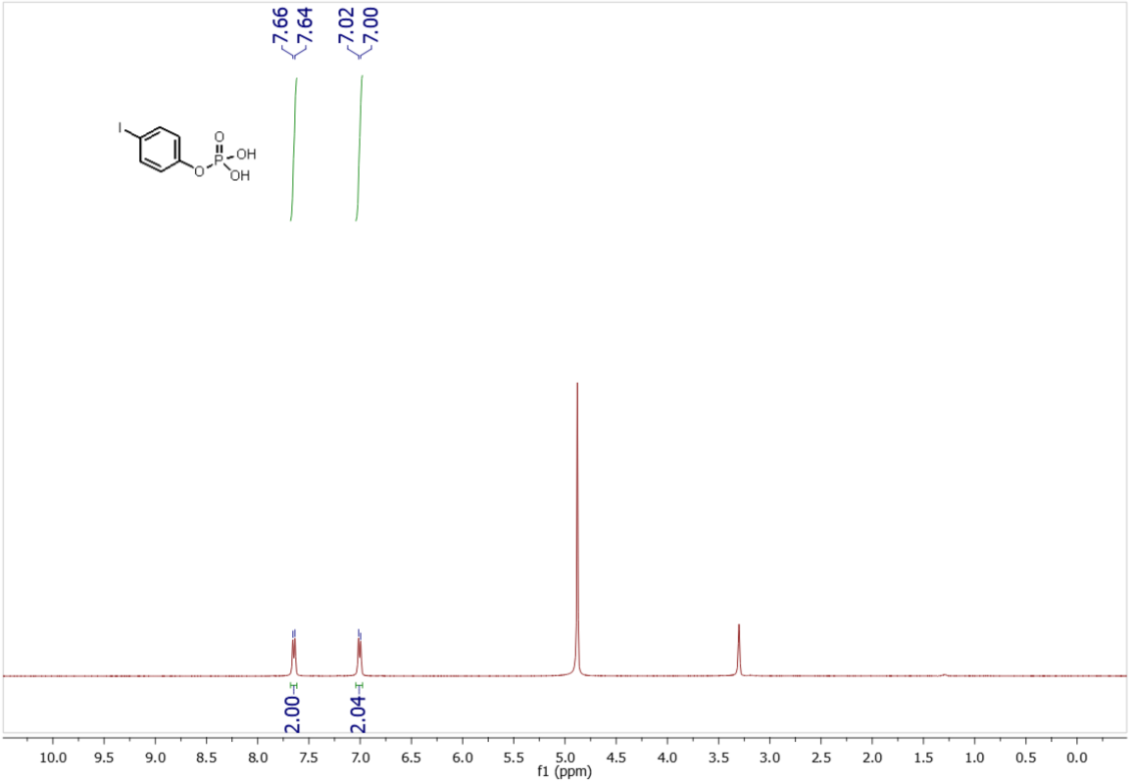


**Figure S23.** $^{1}$H NMR spectrum of 4-iodophenyl dihydrogen phosphate in MeOD.

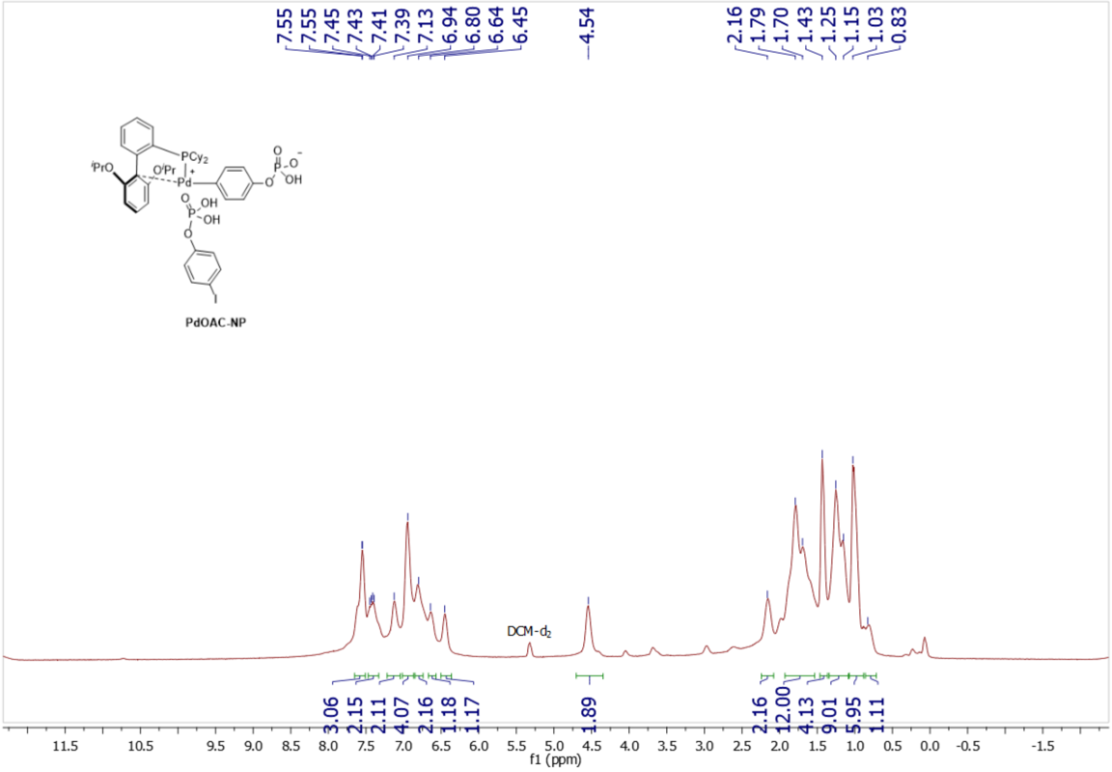


**Figure S24.** $^{1}H$ NMR spectrum of **Pd-OAC-NP** in $CD_2Cl_2$.

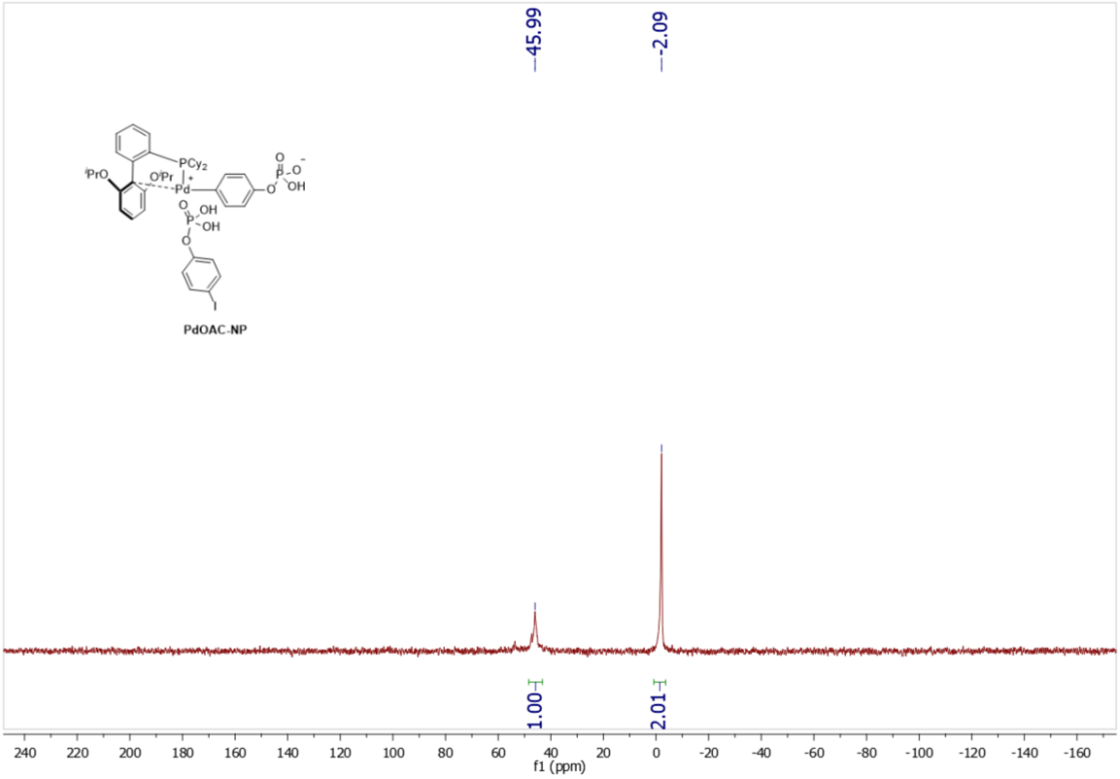


**Figure S25.** $^{31}P$ NMR spectrum of **Pd-OAC-NP** in DMF-$d_7$.

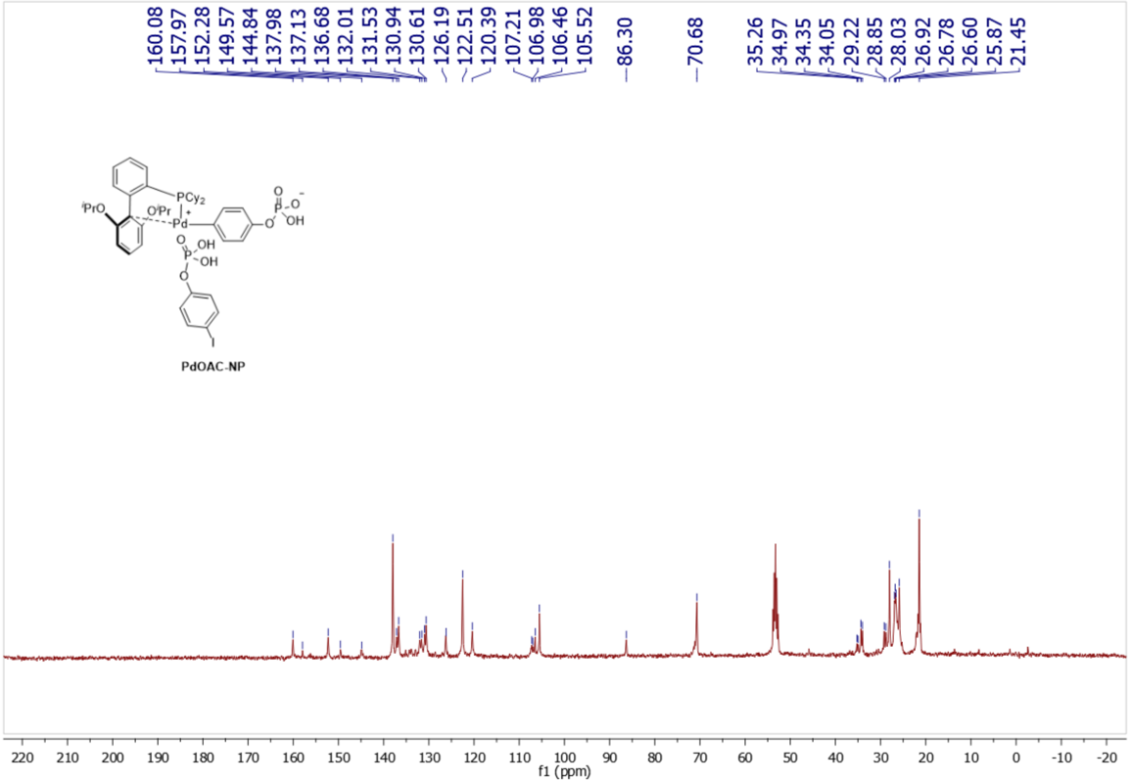


**Figure S26.** [13]C NMR spectrum of **Pd-OAC-NP** in $CD_2Cl_2$.

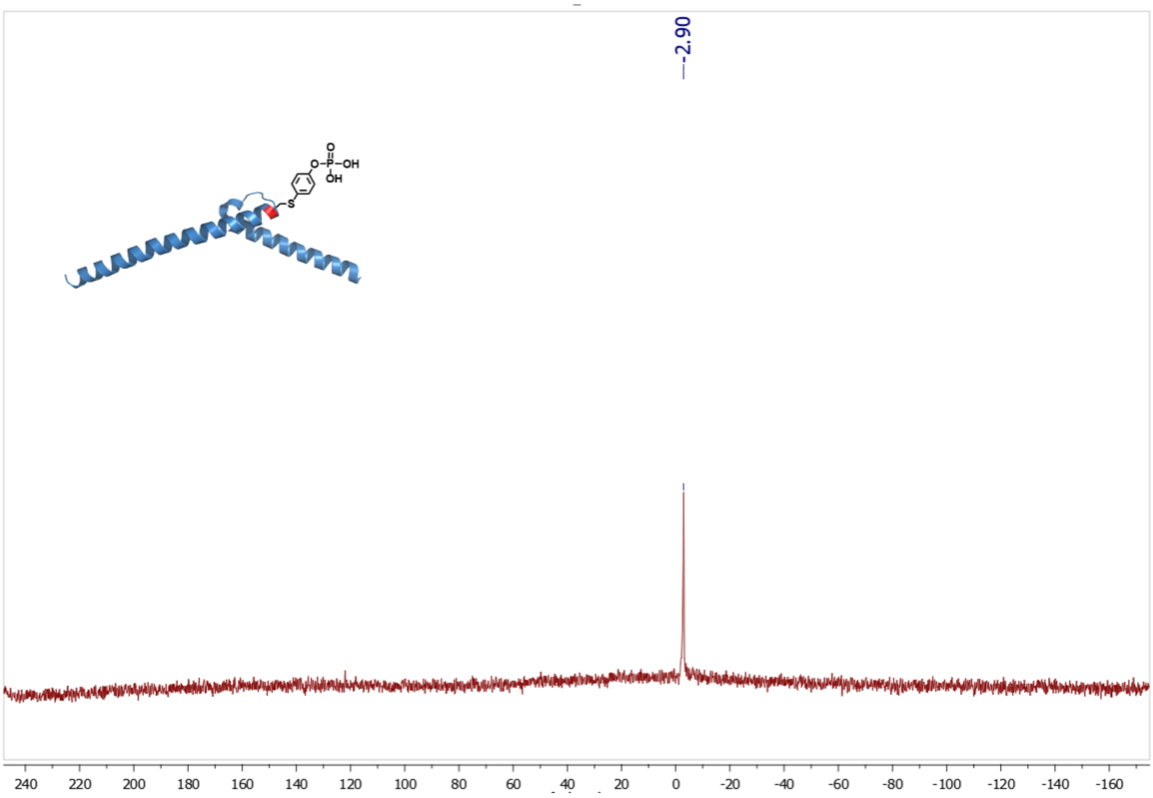


**Figure S27.** [31]P NMR spectrum of **MaxA61C-P(nat)** in $D_2O$.

#### 29. References

1. Miyano, M. Synthesis of organic phosphorus compounds. II. Exhaustive debenzylation reactions. *J. Am. Chem. Soc.* **1955**, *77*, 3524-3526.
2. Vinogradova, E. V.; Zhang, C.; Spokoyny, A. M.; Pentelute B. L.; Buchwald, S. L. Organometallic palladium reagents for cysteine bioconjugation. *Nature* **2015**, *526*, 687-691.
3. Lin, X.; Nithun, R. V.; Samanta, R.; Harel, O.; Jbara, M. Enabling Peptide Ligation at Aromatic Junction Mimics via Native Chemical Ligation and Palladium-Mediated S-Arylation. *Org. Lett.* **2023**, *25*, 4715-4719.
4. Lin, X; Mandal, S.; Nithum, R. V.; Kolla, R.; Bouri, B.; Lashuel, H. A.; Jbara, M. A Versatile Method for Site-Specific Chemical Installation of Aromatic Posttranslational Modification Analogs into Proteins. *J. Am. Chem.Soc*. **2024**, *146*, 37, 25788–25798.
5. Kumar, S. T.; Donzelli, S.; Chiki, A.; Syed, M. M. K.; Lashuel, H. A. A simple, versatile and robust centrifugation-based filtration protocol for the isolation and quantification of α-synuclein monomers, oligomers and fibrils: Towards improving experimental reproducibility in α-synuclein research. *J. Neurochem.* **2020**, *153*, 103–119.
6. Mahul-Mellier, A.-L.; Burtscher, J.; Maharjan, N.; Weerens, L.; Croisier, M.; Kuttler, F.; Leleu, M.; Lashuel, H. A. The process of Lewy body formation, rather than simply α-synuclein fibrillization, is one of the major drivers of neurodegeneration. *Proc. Natl. Acad. Sci. USA* **2020**, *117*, 4971–4982.
